# Cognitive capacity and control in the evolution of intelligence

**DOI:** 10.64898/2026.03.07.710317

**Authors:** Cameron Rouse Turner, Evan M. Russek, Amanda Seed, Emma Suvi McEwen, Natalia Vélez, Thomas J. H. Morgan, Thomas L. Griffiths

**Author notes:** C.T. and E.R. are joint first author contributors. C.T. Conceptualization, Evolutionary modeling, Methodology, Statistical modeling, Writing – original draft. E.R. Conceptualization, Statistical modeling, Methodology, Writing – review and editing. A.S., N.V., T.M., T.G. Conceptualization, Funding acquisition, Writing – review and editing. E.M. Conceptualization, Writing – review and editing. No competing interests to declare.

## Abstract

A diversity of intelligences arises from the constraints under which animals evolve. However, characterizing how constraints shape intelligence is challenging because it requires relating the restrictions on cognitive mechanisms to those that affect their evolution. We demonstrate the potentially complex interaction between constraints by considering the case study of working memory. Here, information-processing capability is limited by the storage *capacity* available to hold representations, and the degree of *control* over those representations. We present an evolutionary model that is mechanistically detailed enough to capture the interactions between capacity and control. This allows us to make quantitative predictions about the distinct patterns of information processing that might be observed across animals. Further, our model’s cognitive detail allows us to fit recall performance on the retro-cue task, illustrating how model predictions can be tested by comparing humans and rhesus monkeys (*Macaca mulatta*). We find that capacity and control are synergistic and amplify each other’s effects. However, evolution prioritizes investment in capacity because it is required for control to be effective. The strength of synergy varies due to interactions between these cognitive components depending on task complexity, cue reliability, and the availability of metabolic energy. Consequently, our model predicts diversity in investment in capacity and control across animals, and identifies a small number of regimes into which lineages could evolve. We discuss how the computational structure of tasks exerts selection on cognitive designs.

**Significance Statement:** Theories about the cognitive processing required for intelligence have been developed largely independently from analyses of evolutionary pressures. This produces an impediment to understanding the diversity of intelligences across animals. We present a mathematical model that bridges these two theoretical domains, focusing on working memory. Our analysis reveals a rich interaction between the capacity to store information and the ability to control that process. We predict that animal intelligences fall into a few major cognitive regimes: initially capacity is prioritized, then capacity and control increase together due to synergy, and finally capacity returns to the fore as adding further control yields diminishing returns. We demonstrate our model makes predictions that can be compared against empirical data via cross-species experiments.

---

**A**key challenge for evolutionary accounts of intelligence is understanding the diverse kinds of intelligence demonstrated across animals. Recent work has argued that intelligence evolves to cope with environmental complexity, which may include ecological circumstances such as unpredictable seasonal fluctuations, or a social life that requires collaboration with group members (1–5). Social and ecological complexity act in concert when species enter a niche that favors both innovation of solutions in response to environmental challenges, as well as acquiring solutions via social learning (6–11). For instance, capuchin monkeys (*Cebinae*) can innovate multistep methods to acquire protected foods (e.g., dehusking Panamá fruit seeds), learn these methods from others, and have unusually large brains (12, 13). Across species, larger brains co-occur with higher rates of innovation and social learning, linking cognitive expansion to the demands of solving new problems (13–15).

While environmental complexity may be the ultimate evolutionary cause of intelligence, it does not explain what *kind* of intelligence should emerge—what cognitive mechanisms should underlie intelligent behavior. An influential line of work from mainstream cognitive science has shown the importance of the information-processing capability of cognitive mechanisms. A central principle provided by this work is that increases to the ability to process information vastly extend the problems a computational system can solve (16–18). This is because when more information can be held in memory to operate on, a greater number of abstract relationships can be identified and used for problem solving. By analogy, when you can see more jigsaw pieces at once, you are able to notice more objects in the scene and successfully put together the puzzle. Therefore, the number of problems a system can solve is bounded by its *capacity*: the upper limit on the amount of information that can be stored, processed, or manipulated. However, the utility of information-processing resources also depends on how well they are managed, meaning that capacity must be considered alongside *control*: the ability to regulate information to achieve goals.

Capacity and control provide different constraints on information-processing capability, and so repeatedly appear as important constructs in explaining performance across cognitive domains. For instance, in the domain of planning, capacity bounds the number of steps an agent can consider, with control being the ability to prioritize favorable sequences and important decision points (19, 20). In visual working memory, capacity is the bound on the details of the scene that can be stored, while control is the ability to internally draw attention to important aspects of the scene (21). Broadly, the degree to which a system can store and manipulate higher-fidelity representations of a task governs its ability to produce solutions (supported formally by the space hierarchy theorem (22)). For example, the ability to detect the repeating pattern in the string ‘AABAAB’ requires being able to hold three symbols. As it cannot be done with two, performance is limited by capacity. Similarly, if representational noise were to corrupt a symbol within the stored ‘AAB’ pattern (risking it becoming ‘AXB’), control processes become crucial to avoid this information loss. Taken together, this suggests a link between information-processing mechanisms and problem solving across taxa. A study of 36 species found a positive association between self-control, brain size, and dietary generality that requires broad problem solving (23). Further, performance across multiple tasks loads onto a single information-processing construct that correlates with brain size across humans, chimpanzees, monkeys, dogs and rodents (24, 25). However, the details of the mechanisms driving these relationships remain elusive (26).

Recently, it has been argued that mechanistic accounts of intelligence based on information processing dovetail with evolutionary explanations that take environmental complexity to be the driver. Highly intelligent animals are under selection to solve the novel problems generated by social and ecological heterogeneity (6–11), while expansions to both capacity and control could equip animals with the cognitive abilities required for success at novel problems (16). For instance, such expansions could support the ability to capture higher-order causal relations, as well as the learning and generalization of complex hierarchical sequences (27–29). The foregrounding of capacity and control makes mechanistically precise the prior suggestion that information processing must expand in response to a higher volume of incoming information from social learning (8–10). However, it leaves open the question of whether these cognitive components would be expected to jointly evolve, whether through synergy or trade-off.

Considering how capacity and control jointly evolve provides a potential path to identifying why different kinds of intelligence might emerge. If there are different adaptive niches in the landscape of capacity and control, we might expect to see lineages diverge into these niches. However, understanding how capacity and control have changed over evolutionary time is a challenge because it requires a detailed understanding of both cognitive mechanisms and evolution. Recent progress appears to have occurred due to efforts to create evolutionary theory that ‘enquires within’, grappling with the details of cognitive operation rather than abstracting over them by taking the ‘phenotypic gambit’ (30–32). A fruitful approach to studying cognition is to consider how a system operates given it must use its limited computational resources to store and manipulate representations as it performs tasks (33). Bringing evolution into view adds a further layer of conceptual difficulty, as we must also consider how evolution divides limited metabolic energy between the structures underpinning cognition over evolutionary time. To be favored, the energetic cost of investing in capacity or control must pay for itself in terms of the fitness gained by an animal. This pressure to balance metabolic investment introduces the possibility of functional trade-offs between capacity and control. Selection works on combinations of traits in the form of whole organisms, so the way in which control processes are facilitated or hindered by available capacity will determine their joint evolution.

Here, we examine the rich interaction between capacity and control, presenting the implications for the evolution of intelligence. We draw on cognitive models of working memory to define an evolutionary model that clarifies the conditions for investment in capacity and control. This approach allows us to find that a rich interaction emerges even in this stylized setting, and thereby make predictions about the diversity of capacity and control across taxa. Consequently, our approach is complementary to the *information capacity hypothesis*, which focuses on improvements to information processing leading to greater problem solving and intelligence (16). We illustrate how our model’s predictions can be empirically validated by comparing working memory performance of humans and rhesus macaques (*Macaca mulatta*). By connecting mechanistic and evolutionary perspectives on cognitive constraints, our work helps explain how the diversity of intelligences seen in nature reflects the availability of resources, and interaction between components of cognition.

## Model

Our model studies selection on an animal who must store a representation of multiple objects in its environment, updating its representation when new information emerges. We build on research into visual working memory and attention, which has benefited from precise formal models and experimental tasks (34–38). We communicate our model through the example of an animal attempting to track the location of fruits during foraging (Fig. 1). The animal experiences similar events as in the well-studied retro-cue task, which is designed to draw on the important functions of working memory (36). The observer must store a representation of multiple items from its visual scene, allocating cognitive resources between items. Representations degrade due to accumulating noise that leads to errors, while control processes actively allocate cognitive resources across items to support recall. If the scene is complex and contains many items, representations erode rapidly. The observer receives a cue indicating which item in memory is likely going to need to be recalled. This cue arrives during retention and acts *retro*actively, providing information that control can use to prioritize the cued item. Using retro-cues requires operations within an internal cognitive workspace that go beyond cues that direct attention while the scene is visible (e.g., sensory filtering). The cue is only partially reliable, meaning uncued items must sometimes be recalled. As correct recall leads to energetic benefits (e.g., fruit), we examine how selection metabolically invests in capacity versus control, and the consequences for information-processing capability.

**Fig. 1.**
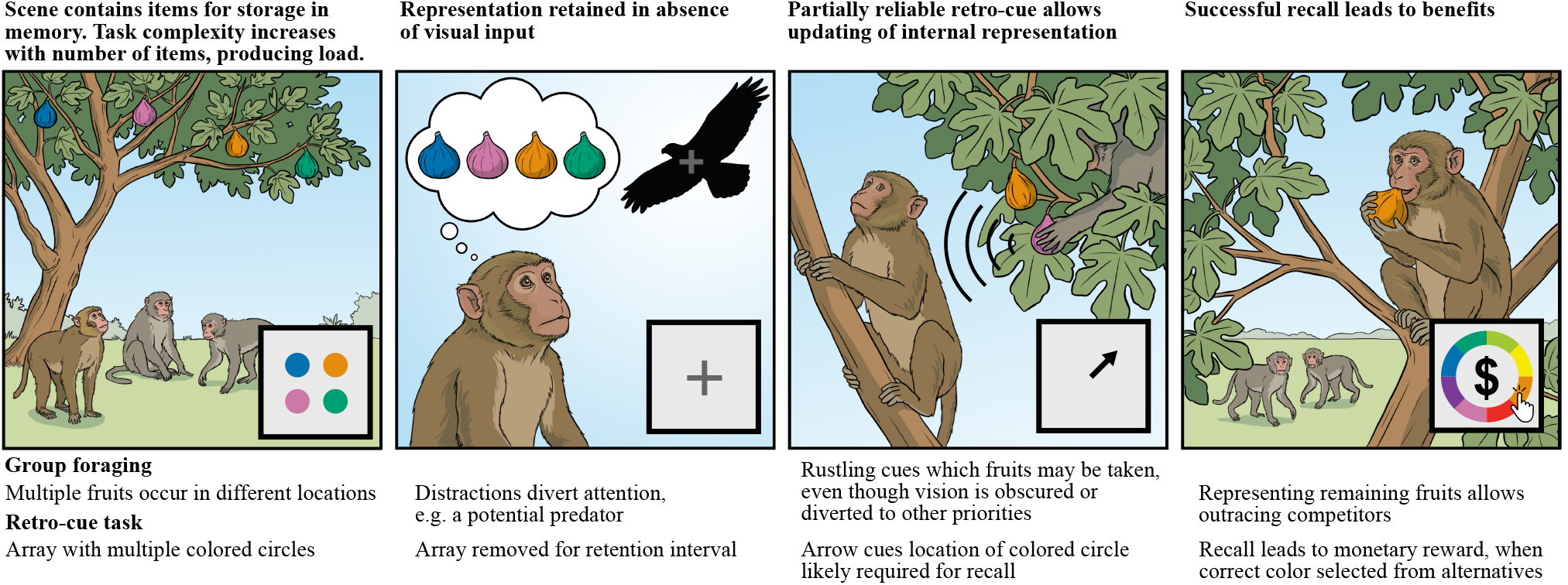
As animals engage in group foraging, working memory operates to overcome similar challenges as the retro-cue task is designed to pose. Panels illustrate natural situations where it is beneficial to encode and manipulate representations of the environment, with insets depicting key screens seen by participants in the retro-cue task. Our model focuses on scenarios where an animal must retain multiple items in memory, because it does not know which item will become useful in the future. Success requires remembering a single item determined externally by the environment, as formalized by *R*(*x*) (Eq. 1). For example, an animal may briefly have the chance to beat competitors to a single remaining fruit whose location must be retrieved from memory. Analogously, in the retro-cue task, the experimenter probes an item whose color the participant must recall.

To make our model tractable, we idealize away realistic details of this scenario. For instance, working memory is particularly useful when a task has not yet become consolidated, but we omit learning dynamics to focus on the selection produced by encountering novel tasks (35). Although in reality capacity and control modulate in response to task demands, we made the simplifying assumption that individuals inherit a fixed capacity and control value that is interpreted as their lifetime expectation. Nonetheless, in the context of models of the evolution of memory, ours is cognitively rich (39–42). Below we outline the model, with detailed derivations in the SI Appendix (Sections 1–3).

### Cognitive processing

Our cognitive model assumes the accuracy of recall results from the allocation of limited cognitive resources among items in working memory. Such resources have been conceived of at a variety of levels of analysis including neural spiking rates, representational signal-to-noise ratios, and bits of encoded information (34, 43–45). We take a broadly compatible approach, simply treating resources as an abstract quantity whose limited availability causes interference and forgetting when spread across multiple items. Capacity is the total amount of resources present, while control is the ability to direct resources to combat noise that corrupts representations (Fig. 2). Our model is supported by forerunners that fit empirical trends in working memory tasks (46–48). We provide a new mathematical continuous-time formulation that allows us to leverage connections to the Ornstein-Uhlenbeck process and optimal control theory (49, 50). The load on memory increases as the number of items that may need to be recalled *m* gets larger. We track the resources allocated to each item by a vector *x* = [*x*_1_, …, *x*_*m*_]^⊤^. The sum of allocations (total resources) is the animal’s overall capacity 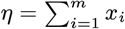. Recall results from how resources are distributed among item representations. The probability of recalling a given item *i* increases with the amount of allocated resources. In particular, we assume that the probability of recall *F* increases monotonically with resource *x*_*i*_. But recall can never be perfect, leading to 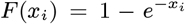. This function captures the commonly arising property of diminishing returns with allocated capacity (43, 51).

**Fig. 2.**
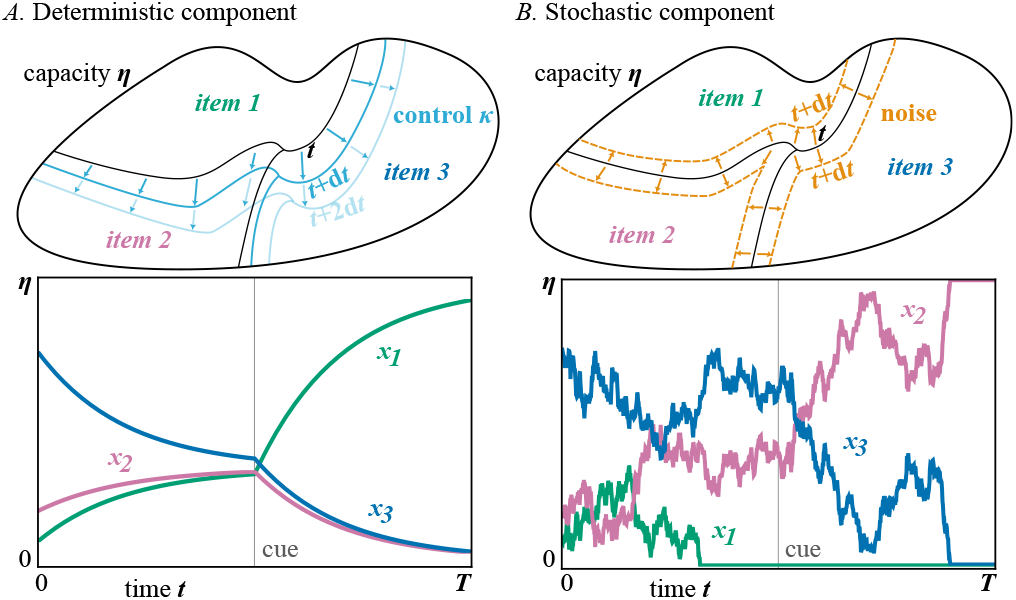
Schematic of deterministic versus stochastic components that combine to give cognitive resource dynamics (SI Appendix, Sections 1–3). Overall capacity is visualized as an abstract volume. We depict storage of three items (*m* = 3), where each item *i* takes up a proportion *x*_*i*_ of the observer’s capacity. **(A)** The deterministic influence of control on resource allocation. Before information is supplied by a cue, control moves resources towards a uniform allocation. After the cue, resources are allocated to the cued item (i.e., item 1). **(B)** The stochastic component of change in resource allocation due to noise, in the absence of control. Noise produces a random change to the assignment of resources, with perturbations drawn from a normal distribution (1*SD* indicated). Our model’s full resource dynamics combine deterministic control (A) with noise (B). Over time, noise causes items that control is trying to protect to erode in memory, resulting in forgetting.

We model a scenario where an observer encounters several items but must later recall only one. The observer does not know which item will become important for recall, although they receive a partially reliable cue indicating the likely target. To calculate an animal’s expected recall, we must consider that cues are not always valid, such that the observer must sometimes recall an uncued item. Letting *q* be the reliability of the cue, the expected recall over cue presentations is:

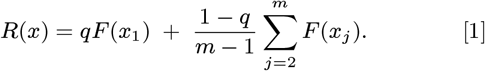

We assign the cued item to be item 1, which has probability *qF* (*x*_1_) of being cued and recalled; otherwise, an uncued item *j* must be recalled each with probability 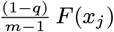. Because *R*(*x*) is concave on our domain, recall improves when control drives the current resource allocation *x* towards an optimal allocation 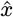. That is, 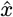 is a desired setpoint that the system is controlled towards (SI Appendix, Sections 1–2). For instance, in the absence of a cue it is optimal to have a uniform allocation 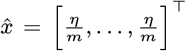. With a perfectly reliable cue, *q* = 1, all resources should be allocated to the cued item 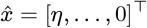.

We assume resources stochastically change in their allocation and thereby corrupt representations due to factors such as noise in neural mechanisms and interference with other tasks. The deterministic action of control and corrupting influence of noise can be modeled by a stochastic differential equation. In particular, for each item *i*, resource dynamics over cognitive processing time *t* are:

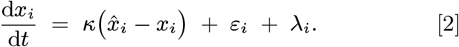

While *R*(*x*) (Eq. 1) is the chance of recall given a resource allocation, d*x*_*i*_*/*d*t* (Eq. 2) governs how allocations change.

We parameterize control as *κ*, the rate of change of the system in a straight line towards the optimum allocation strategy, 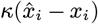. However, each allocation is continuously perturbed by Gaussian white noise *ε*_*i*_(*t*), so the stochastic component of change is driven by a Wiener process. If there is no control *κ* → 0 the observer has no ability to alter resource allocation; as control becomes large *κ* → ∞ resources can be rapidly allocated as the observer dictates. This produces a minimal instantiation of control, encompassing operations that promote task-relevant representations. Capacity *η* is conserved over processing time *t* by subtracting the mean change across allocations 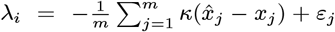. Once an item has no resources allocated it is completely forgotten (when *x*_*i*_ = 0 it remains there at an absorbing boundary). The observer’s strategy changes in response to information provided by the cue. Before the cue, the observer is engaging in a pure recall task attempting to remember all *m* items using the appropriate optimal allocation strategy 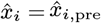 (SI Appendix, Sections 1–3). Receiving an informative cue leads to a change in the optimal allocation strategy 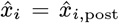. Control then shifts resources, increasing the information the observer’s representation carries about the item they will likely need to recall. Finally, at a uniformly random time within period *T* the observer must engage recall to respond.

### Evolution

Evolution must balance the energy invested in cognition with the benefits it provides. Recall that yields metabolic benefits (e.g., fruit) increases fitness, but at the expense of investment in capacity and control:

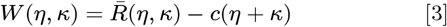

In particular, benefits are given by the expected recall 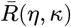, now also taken over noise in cognitive processing and the time of response. Here, *c* is the metabolic cost per unit of investment in cognition relative to the benefits of recall, so weights total investment, *η* + *κ*. Greater capacity is costly because it requires maintaining a reservoir of resources for representation (e.g., more neural activity or larger network). Greater control is costly because it requires mechanisms for effectively routing those resources (e.g., reliable neural gating or stronger modulation of activation). We also examined separate costs for each component. Our model entails no frequency dependence or genetic constraints, and assumes a constant supply of genetic variation. Therefore, we formulate an evolutionary optimality model where populations follow fitness gradients. The cognitive processing that leads to recall 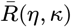 is a complex stochastic process, so we gained an understanding of optimal investment in capacity and control via Monte Carlo simulation and an evolutionary algorithm (SI Appendix, Sections 3–5). Interestingly, our formulation is structurally similar to models capturing the modulation of capacity and control during tasks, and shares broad structure with error-driven learning (48, 52, 53). These models take the optimization objective to be utility rather than lineage fitness.

## Results

### Capacity and control synergize

A central motivation for our model is understanding how capacity and control interact, and if this leads to functional trade-offs or synergies that shape their evolution. Disentangling the roles of capacity and control is a challenge for mainstream cognitive science in its own right (18, 46, 54). In building towards our evolutionary predictions, we first consider their relationship to recall, allowing them to vary freely without metabolic cost (Fig. 3 A and B). We find capacity and control are synergistic: an increase in one enhances the efficacy of the other. For any amount of capacity, increasing control leads to improved recall. This means there is no trade-off such that one cognitive trait produces interference that makes the other detrimental. A larger capacity means more resources to maintain representations, with control allowing the directing of resources in ways that support recall (e.g., towards the cued item). Synergy arises because noise has less of an eroding effect on representations in high-capacity systems, thereby increasing the effective strength of control. Although our model is continuous, consider the analogy of a representation held in a discrete 2-bit memory: two noise events can completely corrupt it, whereas a 100-bit memory would require at least 100 noise events. Control attempts to direct resources but is fighting against buffeting noise. When the effect of noise is reduced by higher capacity, the same amount of control produces more effective allocation.

**Fig. 3.**
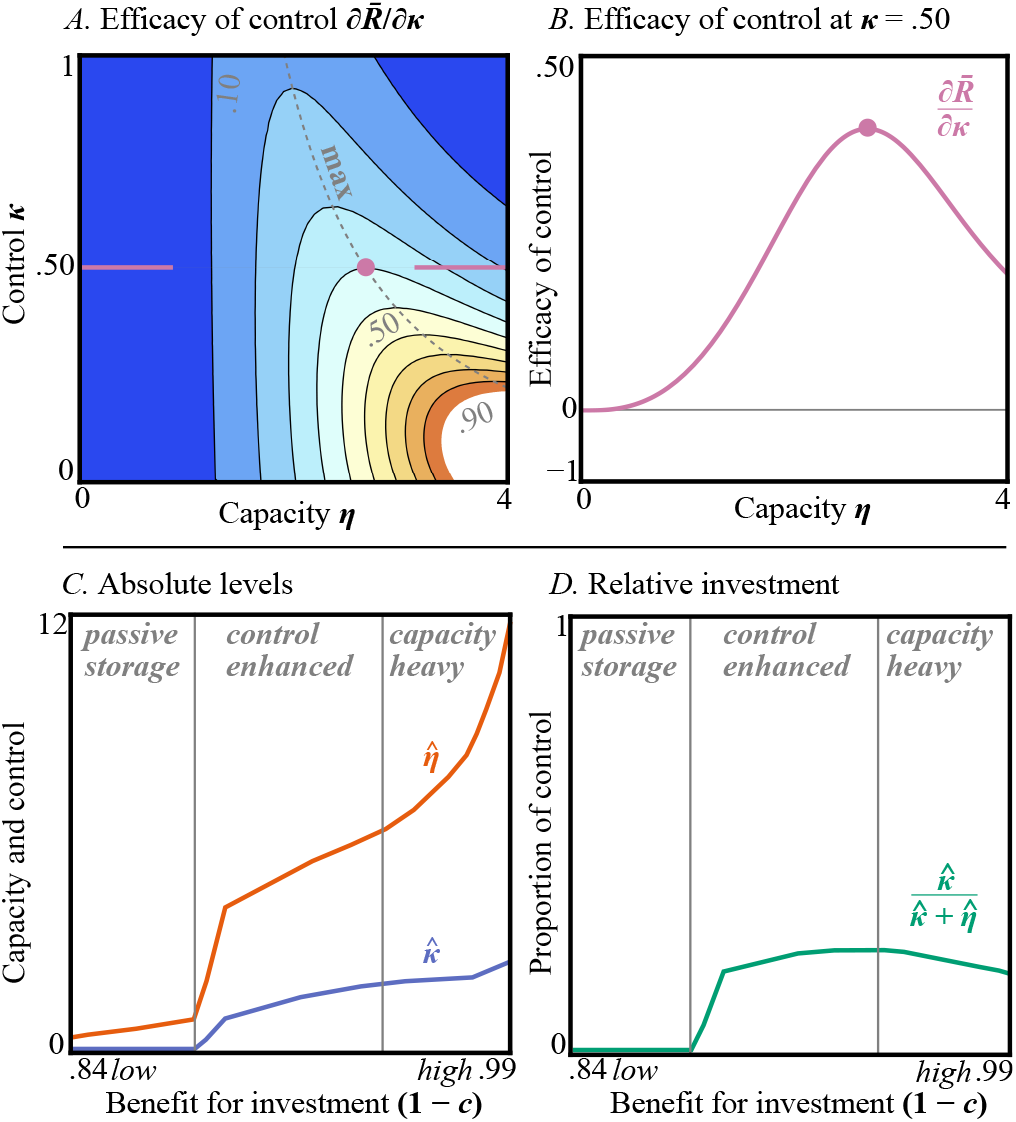
Capacity and control interact to shape recall and evolutionary investment. **(A)** Examining the improvement in recall with an increase in control, 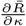, allows us to see how capacity and control moderate each other’s efficiency (see SI Appendix, Sections 6–8). Synergy is present as recall improves with control for all values of capacity (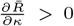 for all *η* and *κ*). Warmer colored contours indicate larger positive values. **(B)** Representative slice of A, showing strongest synergy at intermediate values. **(C)** Evolutionarily optimal values (example sets *m* = 4, *q* = 1, *T* = 3). Maximum metabolic benefits are normalized to be yielded by perfect recall 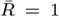; we plot the fraction of maximum returns for a unit of investment, 1 − *c*. Capacity and control both increase when recall leads to higher metabolic benefits, with capacity receiving greater investment. **(D)** Portfolio of investment. At low metabolic benefits pure capacity systems are favored. Moderate benefits allow control-boosted recall. High metabolic benefits lead to heavily capacity-based cognition. Although control processes continue to be enhanced, there is substantial relative investment in capacity. These three regimes emerge because capacity is prioritized, and because of strong synergy at intermediate values (note correspondence of form to B). Figure shows smoothed solutions, for underlying evolutionary simulation output see Fig. S1.

The strongest synergy occurs at intermediate values of capacity and control (Fig. 3B). This is because with too little capacity, control cannot provide much help as noise dominates; whereas with substantial capacity, weak control is already effective. A subtle finding of our model is thus that increasing capacity has a non-monotonic effect on the degree to which control is beneficial.

### Capacity investment prioritized in joint evolution

Our model makes predictions about the pattern of capacity and control across species. We report the optimal capacity-control investment as a function of the metabolic benefit generated by cognition relative to the per-unit cost of investment (Fig. 3 C and D). Broadly, when successful recall leads to substantial benefits (e.g., highly-caloric fruit) this can be expended on producing a sophisticated brain, so evolution favors higher levels of both capacity and control. However, relative investment varies, with evolution prioritizing capacity across all amounts of metabolic benefit. This is because recall can be achieved if there is sufficient capacity, but control cannot operate without a space in which to maintain representations. However, control is still highly valued by evolution, with the fittest cognitively-sophisticated systems investing in both components.

Capacity prioritization should be found across many models where capacity is conceived of as the total available resources, with control being the strength of the deterministic drift that fights diffusive noise. Our mathematical analysis shows that expanding the space of resources broadly produces much greater gains than bolstering the corrective pull (SI Appendix, Sections 7–9). Formally, with regard to the ability to concentrate resources around the optimal allocation, capacity contributes linearly *η*, whereas control suffers severe diminishing returns because it contributes sublinearly 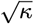. We assume a simple exchange rate for investment in capacity and control, set by the metabolic cost parameter *c*. We also examine an extension with separate costs. Capacity prioritization broadly holds, and this prediction should only be reversed if control is substantially cheaper.

### Information-processing niches

We can identify three regimes of investment in capacity and control that species could occupy (Fig. 3 C and D; SI Appendix, Sections 7–8). When the metabolic benefits achievable by cognition are low, we predict a *passive-storage regime*, in which there is capacity in the absence of control. For species whose cognition generates intermediate levels of benefit, investing in control becomes highly beneficial as it synergizes with capacity, leading to a *control-enhanced regime*. As we consider species for whom cognition generates high metabolic benefits, synergy continues to operate so the value of both capacity and control continue to increase in absolute terms. However, in relative terms, the value of investing in capacity is higher. This leads species with energetically-expensive nervous systems to enter a *capacity-heavy regime*. That is, noise is substantially reduced by high capacity, so control is already efficacious, leading to low marginal gains for additional control investment. A productive way of understanding the diversity of animal intelligences is by considering cognitive transitions over evolutionary history (55–59). Our three regimes suggest a progression over evolutionary time (e.g., capacity first evolves setting the stage for control to be favored). How frequently species occupy these different theoretical regimes in nature remains to be determined.

### Connecting the model to behavioral data

The richness of the description of cognition within our evolutionary model allows us to estimate capacity and control from the behavior of different species. To illustrate how our predictions can be validated, we took advantage of existing data on the performance of 10 monkeys on the retro-cue task (60), adapting the design to run a new experiment with 346 human participants (SI Appendix, Sections 10–13). Stimulus appearance and presentation times replicated those given to monkeys, with adjustments to allow responding by humans online using keyboard and mouse instead of a touchscreen. During a trial, participants had to retain up to four images in memory (Fig. 4). This was followed by a partially reliable retro-cue indicating which image is likely to be required for recall by flashing at its spatial location. Load *m* was manipulated to be two, three, or four image items. Cue reliability *q* was manipulated to be .70 or .92. Pre-cue retention interval was manipulated to be 300 ms, 700 ms, or 1100 ms, with corresponding changes in post-cue interval such that their sum was 1350 ms. A participant’s recall of the original array was either correct or incorrect, allowing us to approximate expected recall over trials. Human participants undertook a single 120-trial session. The monkeys’ sessions were identical in length and they did up to 48 sessions. The human study protocol was approved by the Princeton University Institutional Review Board. All participants provided informed consent online before beginning the task.

**Fig. 4.**
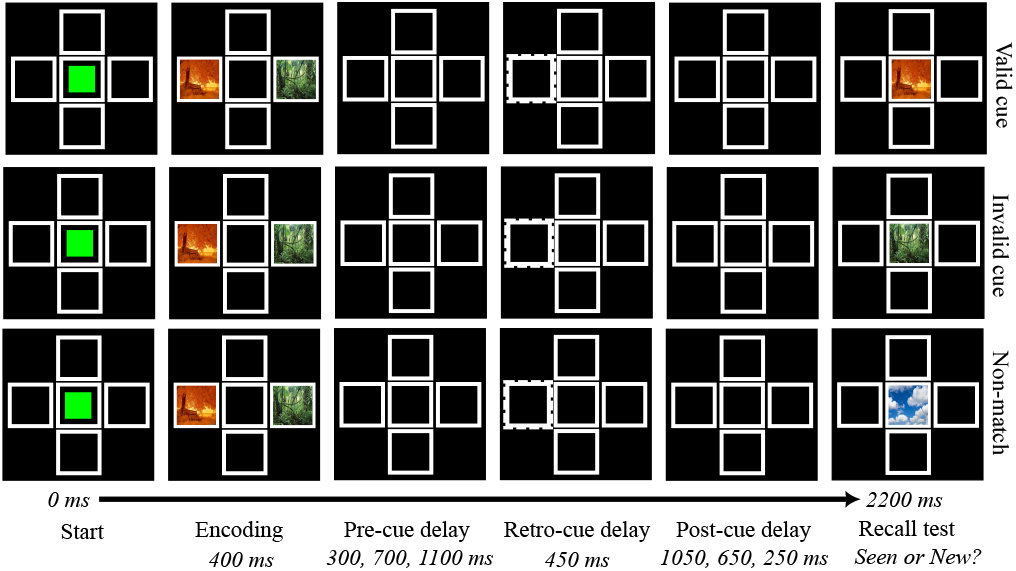
Trial structure for the retro-cue task undertaken by humans and rhesus monkeys. During the encoding, observers saw up to four images of natural scenes in the exterior squares (two depicted). After retention, a flashing retro-cue appeared at one of these locations. The observer’s recall was tested by asking if the image was present during encoding. The cue was partially reliable. Depending on trial-type condition, the cue either validly or invalidly indicated the image to be recalled. Together, valid and invalid cues comprised a total of 60 trials for which the correct response was ‘Seen’. The cue could also be irrelevant, so there were a further 60 non-match trials where the correct response was ‘New’. These trial types were presented randomly within-participants. Load and pre-cue delay were manipulated in a mixed design, being presented in randomized blocks when within-participants.

We implemented the cognitive processing part of our evolutionary model as a hierarchical statistical model using *Stan* (61), fitting it to human and monkey recall data to gain direct estimates of capacity and control for each species (SI Appendix, Sections 11–13). We took advantage of a recently pioneered technique by using a neural network surrogate to produce a deterministic likelihood function that approximates the expectation of our stochastic simulation (62). We verified that our model could explain qualitative trends in human and rhesus monkey memory performance (SI Appendix, Section 12).

We found that humans displayed both higher capacity *η* and control *κ* than rhesus macaques, such that their overall investment in working memory was greater (Fig. 5). This matches model predictions, given humans invest more heavily in their brains, and occupy a complex environment due to cumulative culture (6–11). It also aligns with previous work comparing these species using a different capacity measure (63). Species differences were further supported by a model comparison that found the best performing model assumed humans and monkeys differed in both capacity and control (SI Appendix, Section 14). In addition, we found that capacity values exceeded control for both species (Fig. 5). This is consistent with our model’s prediction that expanding the amount of resources should be prioritized over increasing the rate at which resources are allocated.

**Fig. 5.**
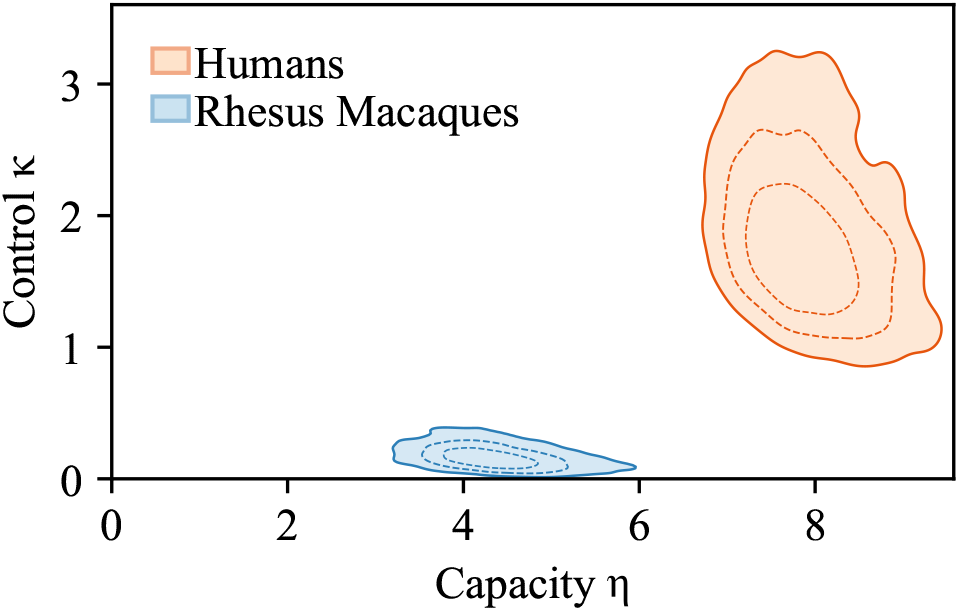
Capacity and control for humans and rhesus macaque monkeys. Shows highest density intervals (HDI) of joint posterior of capacity and control at 50%, 80%, and 95% in expanding contours. Humans have higher joint values of capacity and control than monkeys. Species contrasts show humans have superior capacity to monkeys (M = 3.59, 95% CrI[2.30, 4.93]), and control (M = 1.67, 95% CrI[0.93, 2.80]). Humans invest more in capacity than control (M = 6.07, 95% CrI[4.51, 7.70]), as do monkeys (M = 4.16, 95% CrI[3.25, 5.26]).

Using these behavioral data we provide an example of how to find empirical signatures of our three theoretical regimes for capacity and control. The regimes predicted by our model are identified by the relative investment in control, compared to total investment including capacity, *κ/*(*κ* + *η*) (Fig. 3 D). Both species demonstrate control by using a retro-cue, so they are outside the passive-storage regime, and at least occupy the control-enhanced regime. Humans invest more in aggregate than monkeys. Consequently, if the human proportion of investment in control is also greater, humans are positioned closer to the upper extreme of the control-enhanced regime. Alternatively, if the human proportion of control is lower, they have entered the capacity-heavy regime.

These behavioral data appear consistent with both humans and rhesus monkeys falling within the control-enhanced regime (human proportion control, M = 0.31, 95% CrI[0.24, 0.39]; monkey proportion control, M = 0.05, 95% CrI[0.01, 0.17]; human-monkey difference, M = 0.26, 95% CrI[0.13, 0.35]; SI Appendix, Fig. S11). This is intriguing, given that humans invest substantially in their brain, we might have predicted they reside in the capacity-heavy regime. The central principle of the information capacity hypothesis is that increasing capacity vastly expands task performance (16–18). This seems to provide theoretical backing to the notion that capacity should be disproportionately favored under high investment. By contrast, our results on their face suggest that neither species have entered a regime where increases to control have hit severe diminishing returns. However, we note limitations that suggest caveats to this interpretation. The data reported here are limited in that the procedure was designed specifically for monkeys, and did not push humans close to their extremes (e.g., with a maximum load of 4 items). Further, monkeys received extensive training (48 sessions compared to 1 session for humans). Our intention here is to illustrate an approach to detecting capacity and control, so further research using tailored designs is required to arbitrate between these hypotheses. Recent work has highlighted that careful experiments are required to isolate differences in memory between species from confounds arising from perception or methods of response (64, 65).

## Discussion

The breadth of problems a species can solve is related to the evolution of its capabilities for processing information. While this intuition permeates theory about animal intelligences, little work has formally examined how it connects to contemporary research about processing in cognitive systems. Our approach used a model of the evolution of capacity and control, which are recurring constructs in explanations of the constraints on computational capabilities. We produced a minimal instantiation, focusing on the case of working memory. Capacity was defined as the availability of resources to store a representation of items from the environment, with control being a force that counteracts internal noise to allocate resources using incoming information. Even in our stylized formulation, we showed that there is a rich interaction between capacity and control, with multiple distinct regimes being favored under different environmental conditions. We now consider how the lessons learned in this case study might generalize to the evolution of intelligence more broadly.

### Taxonomizing diversity in capacity and control

Constraints arising from the interaction of cognitive components shape the variety of forms of intelligence that have evolved across animals. We find that capacity and control synergize, improving each other’s efficacy, so cognitively sophisticated species invest in both cognitive components. However, evolution prioritizes relative investment in capacity over control. This occurs in our model because the effects of control are contingent on preexisting capacity.

The varying strength of synergy between capacity and control leads our model to predict that species may evolve into three regimes with regard to the makeup of their information-processing capabilities. Unlike prior verbal taxonomies (56, 57), we give mathematically precise definitions of capacity and control along with quantitative predictions to identify each regime (Fig. 3 C and D). This is achieved by abstracting over different types of control processes (e.g., distractor suppression, representation protection). Nonetheless, the major transitions identified in the control of information over brain evolution broadly cohere with our three regimes (58, 59).

We now speculate about the placement of a few major animal groups, pending detailed empirical work exploring these placements. First, the *passive-storage regime* that arises when there is low investment in working memory appears to approximately describe cognitively basal animals such as jellyfish and sea anemones. The nervous system of these taxa is a simple network with no brain, so while they display short-term storage via sensitization, they lack control functions such as selective attention or shielding from interference (66, 67). Nematodes, flatworms and tardigrades are harder to place. While their nervous systems are rudimentary, they have a centralized controller in the form of a brain-like ganglion (68). These taxa can track context using multiple cues, gating this information to allow integration with memory (69). They appear to be able to shield representations from decay, but manipulating representations within memory using a retro-cue may be out of reach.

Animals that invest heavily in cognition with large brains, such as corvids, elephants, cetaceans, and apes, are predicted to be at least within the *control-enhanced regime*. In these taxa, there is evidence for working memory control processes such as rehearsal, resistance to interference and control of attention (26, 60, 70, 71). Our model predicts that given continued investment, a lineage will eventually hit diminishing returns for control and enter a *capacity-heavy regime*. The principle that increasing information storage vastly expands problem solving appears to support the notion that cognitively sophisticated animals should more strongly prioritize capacity investment than control (16–18). By contrast, our behavioral data suggest rhesus monkeys and humans have a relative proportion of investment in control that locates them both in the control-enhanced regime. Control processes can be more varied and powerful than in our stylized model (72), so there may be extra value that keeps many lineages continually investing in control. Forming strong conclusions about the mapping of taxa to regimes will require further tailored experiments. We hope that such experiments can be facilitated by our efforts to provide a precise and testable understanding of the diversity of information-processing capabilities across diverse lineages.

### Complex tasks and complex minds

Highly intelligent animals succeed by solving complex problems within their environment, an ability that could be underpinned by expansions to capabilities for storing and manipulating information. Our model points to elaborations of this *information capacity hypothesis* (16) by delineating how selection acts on capacity and control given their role in solving ecological challenges. We focused on the task of tracking multiple items, which occurs in many contexts, such as group foraging where the location of food is monitored. However, tasks in nature vary in ways that may create different demands on processing, such as in hierarchical structure (27–29), or the requirement for integration across modalities (73). Even within multi-item tracking, task compressibility depends on item arrangement and distractors, with cognition able to respond by chunking visual regularities to make effective use of limited capacity (38, 74). Future research should examine how aspects of task structure shape the evolution of cognitive design.

Our model suggests superior information-processing capabilities are favored in response to complex tasks if there is reliable information. In the SI Appendix (Section 15), we explore a way to summarize task complexity and map it to cognitive ability using information theory. Although tasks and minds come in many forms, we propose a coherent approach that could be applied across animals. We formally define *task complexity* as the number of bits required to encode the solution. For instance, in our memory task with 4 items, the complexity is 2 bits. Further, we define the observer’s *information-processing capability* to be the information contained in the task weighted by their performance. Consistent with a variety of prior models, we find cognitive abilities for processing information are favored when cues are informative (1, 2, 40–42, 75–78). This is because while complex tasks demand capacity for storage, reliable cues provide a role for control to implement information to allow success. Furthermore, we apply our measures to empirical data, finding that humans’ information-processing capability is superior to that of rhesus monkeys (Fig. S13).

### Limitations and future directions

Explaining the diversity of animal intelligences requires understanding how constraints shape both the breadth and particular kinds of problems a species can solve. Our model focuses on broad information processing via operations in working memory, and thus foregrounds general capabilities for solving novel problems. However, animal cognition is not solely content-neutral, but instead contains structure and biases that aid in solving the idiosyncratic problems faced by that species (79–82). For instance, monkeys rapidly detect snakes, supporting a system for avoiding this threat (83). Understanding how to relate domain-specific and domain-general cognition remains a crucial challenge in the study of diverse intelligences, where we are interested in the scope of the problems taxa can solve. Recent theory argues that greater cognitive specialization may occur when information capacity is limited, because it becomes difficult to store distinct representations for different tasks (84). This suggests an extension of our model considering multiple tasks that require partially overlapping information. As information-processing capabilities are restricted, specializing on a subset of ways of being intelligent may be adaptive. Furthermore, specialization is thought to often occur by the evolutionary canalization of abilities that are initially constructed using domain-general capabilities (82, 85). A complicating factor requiring clarification is that such capabilities themselves may be partially learned, with capacity and control improving over development (46, 54). Longitudinal measurements of working memory suggest age-related changes in capacity and control are partly driven by brain maturation (86, 87). To disentangle the contribution of learning, the features of tasks encountered over development must be measured or manipulated to examine their influence on downstream capacity and control, analogous to studies of early enrichment in monkeys and rodents (88–90). Finally, we predict that capacity and control vary depending on the availability of metabolic energy, task complexity, and informativeness of cues. This highlights that diversity in animal cognition could arise from differences in domain-general abilities.

### Conclusion

Selection hones animal minds but its action is shaped by idiosyncratic constraints, together this guides lineages into diverse forms of intelligence. However, the notion of constraint is a placeholder for a plethora of factors that shape the direction of evolution. Following the present model, we are in a better position to form a principled understanding of the broad types of constraint acting on the evolution of intelligence (84, 91–93). We found an important class of constraints is the availability of resources required for performance. Within cognitive processing, working memory capacity is a limited resource that constrains storage. At the evolutionary level, metabolic energy is a limited resource that must be judiciously invested in different cognitive components to improve fitness. The availability of energy ultimately bounds the sophistication of the brain and cognition. A second broad class of constraints comprises functional tradeoffs or synergies emerging from the interaction of components. These functional interactions govern the performance of the overall system. We found that capacity and control interacted such that the strength of synergy varied depending on level of investment, generating the possibility of different regimes lineages may enter. Only by creating theory that connects constraints on cognition with their impact on the course of evolution can we understand the variety across animal minds.

## Supporting Information Appendix (SI)

SI Appendix available online. Code and data available in our [repository](https://github.com/evanrussek/Capacity_and_Control_Experiment_Analysis).

## ACKNOWLEDGMENTS

We thank Ryan J. Brady and Robert R. Hampton for providing data and materials from their rhesus monkey experiments. We thank Stephen Mann for comments and the Templeton World Charity Foundation for funding (grant number 20648). All AI-assisted output was supervised and reviewed by the authors, who take full responsibility for the final manuscript. Generative AI tools Claude (Opus 4.7), ChatGPT (5.5), and Gemini (3.1) were used to assist with code generation, proofreading and editing, and literature searches. We thank Luis Mora for illustrating Fig. 1.

## References

1. P Godfrey-Smith, Complexity and the Function of Mind in Nature. (Cambridge University Press, Cambridge), (1996).

2. CR Turner, TJH Morgan, TL Griffiths, Environmental complexity and regularity shape the evolution of cognition. Proc. Royal Soc. B: Biol. Sci. 291, 20241524 (2024).

3. S Pinker, The cognitive niche: coevolution of intelligence, sociality, and language. Proc. Natl. Acad. Sci. United States Am. 107, 8993–8999 (2010).

4. D Sol, Revisiting the cognitive buffer hypothesis for the evolution of large brains. Biol. Lett. 5, 130–133 (2009).

5. K Sterelny, Thought in a Hostile World: The Evolution of Human Cognition. (Blackwell Publishing, Oxford), (2003).

6. R Boyd, PJ Richerson, J Henrich, The cultural niche: why social learning is essential for human adaptation. Proc. Natl. Acad. Sci. United States Am. 108, 10918–10925 (2011).

7. E Herrmann, J Call, MV Hernández-Lloreda, B Hare, M Tomasello, Humans have evolved specialized skills of social cognition: The cultural intelligence hypothesis. Science 317, 1360–1366 (2007).

8. J Henrich, The Secret of Our Success: How Culture is Driving Human Evolution, Domesticating Our Species, and Making Us Smarter. (Princeton University Press, Princeton), (2015).

9. C Heyes, Cognitive Gadgets: The Cultural Evolution of Thinking. (Harvard University Press, Cambridge, MA), (2018).

10. KN Laland, Darwin’s Unfinished Symphony: How Culture Made the Human Mind. (Princeton University Press, Princeton), (2017).

11. K Laland, A Seed, Understanding human cognitive uniqueness. Annu. Rev. Psychol. 72, 689–716 (2021).

12. BJ Barrett, RL McElreath, SE Perry, Pay-off-biased social learning underlies the diffusion of novel extractive foraging traditions in a wild primate. Proc. Royal Soc. B: Biol. Sci. 284, 20170358 (2017).

13. SE Street, AF Navarrete, SM Reader, KN Laland, Coevolution of cultural intelligence, extended life history, sociality, and brain size in primates. Proc. Natl. Acad. Sci. 114, 7908–7914 (2017).

14. M González-Forero, A Gardner, Inference of ecological and social drivers of human brain-size evolution. Nature 557, 554–557 (2018).

15. L Lefebvre, SM Reader, D Sol, Brains, innovations and evolution in birds and primates. Brain, Behav. Evol. 63, 233–246 (2004).

16. JF Cantlon, S. Piantadosi, Uniquely human intelligence arose from expanded information capacity. Nat. Rev. Psychol. 3, 275–293 (2024).

17. GS Halford, WH Wilson, S Phillips, Processing capacity defined by relational complexity: implications for comparative, developmental, and cognitive psychology. Behav. Brain Sci. 21, 803–831 (1998).

18. S Musslick, JD Cohen, Rationalizing constraints on the capacity for cognitive control. Trends Cogn. Sci. 25, 757–775 (2021).

19. I Kuperwajs, EM Russek, MG Mattar, WJ Ma, TL Griffiths, Looking deeper into the algorithms underlying human planning. Trends Cogn. Sci. (2025).

20. P Fernandez Velasco, et al., Expert navigators deploy rational complexity–based decision precaching for large-scale real-world planning. Proc. Natl. Acad. Sci. 122, e2407814122 (2025).

21. A Gazzaley, AC Nobre, Top-down modulation: bridging selective attention and working memory. Trends Cogn. Sci. 16, 129–135 (2012).

22. CH Papadimitriou, Computational Complexity. (Addison-Wesley, Reading, MA), (1994).

23. EL MacLean, et al., The evolution of self-control. Proc. Natl. Acad. Sci. United States Am. 111, E2140–E2148 (2014).

24. JM Burkart, MN Schubiger, CP van Schaik, The evolution of general intelligence. Behav. Brain Sci. 40, e195 (2017).

25. RW Engle, Working memory and executive attention: A revisit. Perspectives on Psychol. Sci. 13, 190–193 (2018).

26. CJ Völter, B Tinklenberg, J Call, AM Seed, Comparative psychometrics: establishing what differs is central to understanding what evolves. Philos. Transactions Royal Soc. B: Biol. Sci. 373, 20170283 (2018).

27. S Ferrigno, SJ Cheyette, S. Piantadosi, JF Cantlon, Recursive sequence generation in monkeys, children, U.S. adults, and native amazonians. Sci. Adv. 6, eaaz1002 (2020).

28. GS Halford, WH Wilson, S Phillips, Relational knowledge: The foundation of higher cognition. Trends Cogn. Sci. 14, 497–505 (2010).

29. M Hahn, R Futrell, R Levy, E Gibson, A resource-rational model of human processing of recursive linguistic structure. Proc. Natl. Acad. Sci. United States Am. 119, e2122602119 (2022).

30. C Heyes, Enquire within: cultural evolution and cognitive science. Philos. Transactions Royal Soc. B: Biol. Sci. 373, 20170051 (2018).

31. JM McNamara, Game theory in biology: Moving beyond functional accounts. Am. Nat. 199, 179–193 (2022).

32. TJH Morgan, MW Feldman, Human culture is uniquely open-ended rather than uniquely cumulative. Nat. Hum. Behav. 9, 28–42 (2025).

33. F Lieder, TL Griffiths, Resource-rational analysis: Understanding human cognition as the optimal use of limited computational resources. Behav. Brain Sci. 43, e1 (2019).

34. PM Bays, S Schneegans, WJ Ma, TF Brady, Representation and computation in visual working memory. Nat. Hum. Behav. 8, 1016–1034 (2024).

35. RM Shiffrin, Modeling memory and perception. Cogn. Sci. 27, 341–378 (2003).

36. AS Souza, K Oberauer, In search of the focus of attention in working memory: 13 years of the retro-cue effect. Attention, Perception, Psychophys. 78, 1839–1860 (2016).

37. F van Ede, AC Nobre, Turning attention inside out: how working memory serves behavior. Annu. Rev. Psychol. 74, 137–165 (2023).

38. GA Alvarez, SL Franconeri, How many objects can you track? Evidence for a resource-limited attentive tracking mechanism. J. Vis. 7, 14.1–14.10 (2007).

39. M Ammar, L Fogarty, A Kandler, Social learning and memory. Proc. Natl. Acad. Sci. United States Am. 120, e2310033120 (2023).

40. S Dridi, L Lehmann, Environmental complexity favors the evolution of learning. Behav. Ecol. 27, 842–850 (2016).

41. TJH Morgan, JW Suchow, TL Griffiths, Experimental evolutionary simulations of learning, memory and life history. Philos. Transactions Royal Soc. B: Biol. Sci. 375, 20190504 (2020).

42. PM Todd, GF Miller, Exploring adaptive agency II: Simulating the evolution of associative learning in From Animals to Animats, Proceedings of the First International Conference on Simulation of Adaptive Behavior. (MIT Press), pp. 306–315 (1991).

43. S Schneegans, R Taylor, PM Bays, Stochastic sampling provides a unifying account of visual working memory limits. Proc. Natl. Acad. Sci. United States Am. 117, 20959–20968 (2020).

44. MG Stokes, ‘Activity-silent’ working memory in prefrontal cortex: a dynamic coding framework. Trends Cogn. Sci. 19, 394–405 (2015).

45. CR Sims, Rate–distortion theory and human perception. Cognition 152, 181–198 (2016).

46. E Russek, et al., Modeling the contributions of capacity and control to working memory development in Proceedings of the 46th Annual Meeting of the Cognitive Science Society. Vol. 46, pp. 245–251 (2024).

47. JW Suchow, DD Bourgin, TL Griffiths, Evolution in mind: evolutionary dynamics, cognitive processes, and Bayesian inference. Trends Cogn. Sci. 21, 522–530 (2017).

48. R van den Berg, WJ Ma, A resource-rational theory of set size effects in human visual working memory. eLife 7, e34963 (2018).

49. WH Fleming, RW Rishel, Deterministic and Stochastic Optimal Control. (Springer, New York), (1975).

50. CW Gardiner, Handbook of Stochastic Methods. (Springer, Berlin), 3 edition, (2004).

51. JJ Hopfield, Neural networks and physical systems with emergent collective computational abilities. Proc. Natl. Acad. Sci. United States Am. 79, 2554–2558 (1982).

52. A Shenhav, MM Botvinick, JD Cohen, The expected value of control: An integrative theory of anterior cingulate cortex function. Neuron 79, 217–240 (2013).

53. A Luzardo, E Alonso, E Mondragón, A Rescorla-Wagner drift-diffusion model of conditioning and timing. PLoS Comput. Biol. 13, e1005796 (2017).

54. A Shimi, G Scerif, Towards an integrative model of visual short-term memory maintenance: evidence from the effects of attentional control, load, decay, and their interactions in childhood. Cognition 169, 61–83 (2017).

55. P Bateson, KN Laland, Tinbergen’s four questions: an appreciation and an update. Trends Ecol. Evol. 28, 712–718 (2013).

56. P Carruthers, Evolution of working memory. Proc. Natl. Acad. Sci. 110, 10371–10378 (2013).

57. P Cisek, Evolution of behavioural control from chordates to primates. Philos. Transactions Royal Soc. B: Biol. Sci. 377, 20200522 (2022).

58. AB Barron, M Halina, C Klein, Transitions in cognitive evolution. Proc. Royal Soc. B: Biol. Sci. 290, 20230671 (2023).

59. S Coombs, M Trestman, A multi-trait embodied framework for the evolution of brains and cognition across animal phyla. Behav. Brain Sci. 48, e77 (2025).

60. RJ Brady, RR Hampton, Post-encoding control of working memory enhances processing of relevant information in rhesus monkeys (Macaca mulatta). Cognition 175, 26–35 (2018).

61. Stan Development Team, Stan reference manual (2024).

62. A Fengler, LN Govindarajan, T Chen, MJ Frank, Likelihood approximation networks (LANs) for fast inference of simulation models in cognitive neuroscience. eLife 10, e65074 (2021).

63. LC Elmore, et al., Visual short-term memory compared in rhesus monkeys and humans. Curr. Biol. 21, 975–979 (2011).

64. S Ghirlanda, J Lind, M Enquist, Memory for stimulus sequences: A divide between humans and other animals? Royal Soc. Open Sci. 4, 161011 (2017).

65. E Reindl, AM Seed, RA Barton, T Francis-Costa, RL Kendal, Humans may not have a uniquely enhanced sequence memory: sequence discrimination is facilitated by causal-logical framing in humans and chimpanzees. Royal Soc. Open Sci. 12, 250236 (2025).

66. K Cheng, Learning in Cnidaria: A systematic review. Learn. Behav. 49, 175–189 (2021).

67. T Lev-Ari, H Beeri, Y Gutfreund, The ecological view of selective attention. Front. Integr. Neurosci. 16, 856207 (2022).

68. P Sterling, S Laughlin, Principles of Neural Design. (MIT Press, Cambridge, MA), (2015).

69. JA Haley, SH Chalasani, C. elegans foraging as a model for understanding the neuronal basis of decision-making. Cell. Mol. Life Sci. 81, 252 (2024).

70. RJ Brady, RR Hampton, Nonverbal working memory for novel images in rhesus monkeys. Curr. Biol. 28, 3903–3910.e3 (2018).

71. E Fongaro, J Rose, Crows control working memory before and after stimulus encoding. Sci. Reports 10, 3253 (2020).

72. D Badre, Cognitive control. Annu. Rev. Psychol. 76, 167–195 (2025).

73. AD Baddeley, The episodic buffer: A new component of working memory? Trends Cogn. Sci. 4, 417–423 (2000).

74. TF Brady, T Konkle, GA Alvarez, Compression in visual working memory: Using statistical regularities to form more efficient memory representations. J. Exp. Psychol. Gen. 138, 487–502 (2009).

75. K Aoki, MW Feldman, Evolution of learning strategies in temporally and spatially variable environments: a review of theory. Theor. Popul. Biol. 91, 3–19 (2014).

76. MC Donaldson-Matasci, CT Bergstrom, M Lachmann, The fitness value of information. Oikos 119, 219–230 (2010).

77. AS Dunlap, DW Stephens, Reliability, uncertainty, and costs in the evolution of animal learning. Curr. Opin. Behav. Sci. 12, 73–79 (2016).

78. CR Turner, TJH Morgan, TL Griffiths, Complex brains allow functioning in a complex environment by using information. Behav. Brain Sci. 48, e96 (2025).

79. TL Griffiths, N Chater, C Kemp, A Perfors, JB Tenenbaum, Probabilistic models of cognition: exploring representations and inductive biases. Trends Cogn. Sci. 14, 357–364 (2010).

80. MA Krause, Evolutionary perspectives on learning: conceptual and methodological issues in the study of adaptive specializations. Animal Cogn. 18, 807–820 (2015).

81. PM Todd, R Hertwig, U Hoffrage, Evolutionary cognitive psychology in The Handbook of Evolutionary Psychology, ed. D Buss. (Wiley, Hoboken, NJ), (2016).

82. CR Turner, LD Walmsley, Preparedness in cultural learning. Synthese (2020).

83. A Öhman, S Mineka, Fears, phobias, and preparedness: Toward an evolved module of fear and fear learning. Psychol. Rev. 108, 483–522 (2001).

84. CR Turner, D Argumugam, L Nelson, T Griffiths, Trade-offs between tasks induced by capacity constraints bound the scope of intelligence in Proceedings of the 47th Annual Meeting of the Cognitive Science Society. Vol. 47, pp. 613–620 (2025).

85. TJH Morgan, JW Suchow, TL Griffiths, What the baldwin effect affects depends on the nature of plasticity. Cognition 197, 104165 (2020).

86. SF Ahmed, A Ellis, KP Ward, N Chaku, PE Davis-Kean, Working memory development from early childhood to adolescence using two nationally representative samples. Dev. Psychol. 58, 1962–1973 (2022).

87. B Tervo-Clemmens, et al., A canonical trajectory of executive function maturation from adolescence to adulthood. Nat. Commun. 14, 6922 (2023).

88. H van Praag, G Kempermann, FH Gage, Neural consequences of environmental enrichment. Nat. Rev. Neurosci. 1, 191–198 (2000).

89. A Massera, et al., Longitudinal effects of early psychosocial deprivation on macaque executive function: Evidence from computational modelling. Proc. Royal Soc. B: Biol. Sci. 290, 20221993 (2023).

90. N Cowan, Working memory maturation: Can we get at the essence of cognitive growth? Perspectives on Psychol. Sci. 11, 239–264 (2016).

91. TJ Garland, CJ Downs, AR Ives, Trade-offs (and constraints) in organismal biology. Physiol. Biochem. Zool. 95, 82–112 (2022).

92. TL Griffiths, Understanding human intelligence through human limitations. Trends Cogn. Sci. 24, 873–883 (2020).

93. C Klein, Mechanisms, resources, and background conditions. Biol. Philos. 33 (2018).

