## Supplementary Information (SI) Appendix for "Cognitive capacity and control in the evolution of intelligence"

**PNAS**

**Turner et al. 2026**

***Guide to Supporting Information***

This *SI Appendix* outlines derivation and results.

All model derivations, plotting, and experimental materials available through our repository.

*Contents*


| | |
|---|---|
| 1 Cognitive resource allocation dynamics | 3 |
| 2 Resource allocation dynamics via optimal control theory | 6 |
| 3 Time course of cognitive processing events | 8 |
| 4 Fitness function | 9 |
| 5 Finding evolutionary optimum | 9 |
| 6 Long-run expected recall approximation | 12 |
| 7 Capacity and control interactions | 14 |
| 8 Metabolic costs and why capacity is more efficacious than control | 16 |
| 9 Scale of capacity and control | 20 |
| 10 Experimental test methods | 21 |
| 11 Experimental test statistical analyses | 26 |
| 12 Experimental test results | 28 |
| 13 Full statistical model and derived quantities | 36 |
| 14 Model comparison of species differences in capacity and control | 40 |
| 15 Complex tasks select for information-processing capability | 42 |
| References | 47 |

### 1 Cognitive resource allocation dynamics

An observer has memory resources distributed among $m$ items so that their resource distribution is $x = [x_1, \dots, x_m]^\top$. The resources allocated to any one item must be positive $x_i \geq 0$, with the sum of allocations being the observer's overall memory capacity $\eta = \sum_{i=1}^{m} x_i$. In particular, $x \in \Omega^m \subset \mathbb{R}^m$, where $\Omega^m$ is a regular $m$-simplex.

The probability of recalling an item $i$ is a function $F$ that increases monotonically with the resources allocated to it $x_i$. However, recall can never be perfect, so there are diminishing returns. That is, $F: [0, \eta] \to [0,1)$, such that $\frac{\mathrm{d}F}{\mathrm{d}x_i} > 0$ and $\frac{\mathrm{d}^2F}{\mathrm{d}x_i^2} < 0$ for all $x_i$. Therefore, we use the convenient form: $F(x_i) = 1 - \mathrm{e}^{-x_i}$.

Given the cued item has probability $q \in [1/m, 1]$ of being required to be recalled, the expected recall over cue presentations is:

$$R(x) = qF(x_1) + \frac{(1-q)}{(m-1)} \sum_{j=2}^{m} F(x_j) \tag{S1}$$

In particular, $R$ is the probability of successful recall of a given single item of the original set, after the retro-cue has been presented, conditional on a resource allocation $x$. Our measure of success $R$ differs from correctly recalling any item at all, or recalling all items; in these scenarios a retro-cue is not useful because no item needs to be prioritized. The special case where only the cued item must be recalled is produced by setting $q = 1$.

There is a single optimum resource allocation $\hat{x} = \underset{x \in \Omega^m}{\operatorname{argmax}} R$. This can be found by using the method of Lagrange, by using the constraint $\eta = \sum_{i=1}^{m} x_i$ and solving $\nabla R = \alpha\nabla[x_1 + \cdots + x_m - \eta]$, where $\alpha$ is the Lagrange multiplier. On our domain, $\Omega^m$, the Hessian of the Lagrangian is negative semidefinite:

$$H[\mathcal{L}(R)] = \begin{bmatrix} -q\mathrm{e}^{-x_1} & 0 & \cdots & 0 \\ 0 & -\frac{(1-q)}{(m-1)}\mathrm{e}^{-x_2} & \cdots & 0 \\ \vdots & \vdots & \ddots & \vdots \\ 0 & 0 & \cdots & -\frac{(1-q)}{(m-1)}\mathrm{e}^{-x_m} \end{bmatrix} \tag{S2}$$

That is, eigenvalues appear along the diagonal, and they are always all negative or zero. Therefore, $R(x)$ is concave subject to our constraints, with a unique global maximum at $\hat{x}$.

The optimal allocation for the cued item 1, and the optimal of any of the other $j$ items, are respectively:

$$\hat{x}_1 = \frac{\eta}{m} + \frac{(m-1)}{m}\ln\left((m-1)\frac{q}{(1-q)}\right) \tag{S3}$$

$$\hat{x}_j = \frac{\eta}{m} - \frac{1}{m}\ln\left((m-1)\frac{q}{(1-q)}\right) \tag{S4}$$

As the reliability of $q$ increases it is optimal to allocate more resources to the cued item. It is also ideal to allocate more resources to the cued item when $m$ increases because the a priori probability of remembering any item goes down, meaning the cue carries more information. As $\eta$ increases it is optimal to have a more uniform distribution of resources; this is due to the diminishing returns of improving the recall. New capacity can improve the recall of the cued item relatively less than the un-cued items. When a cue provides no information ($q = 1/m$) about the item to be recalled, control pushes resources towards a uniform distribution $\hat{x} =$

$\left[\frac{\eta}{m}, \ldots, \frac{\eta}{m}\right]^\top$; this recovers pure retention and the situation before a cue has arrived. When a cue is perfectly reliable ($q = 1$) control pushes all resources towards the cued item $\hat{x} = [\eta, \ldots, 0]^\top$. The optimum resource allocation goal $\hat{x}$ is a desired setpoint that cognitive control pushes resource allocation towards. Given $\hat{x}$ is a global maximizer, expected recall increases by moving $x$ along as straight a path as possible towards $\hat{x}$. This justifies the general form of our resource dynamics equation, in which direct motion towards $\hat{x}$ occurs at a rate given by cognitive control $\kappa \in [0, \infty)$, under the constraint that capacity $\eta \in (0, \infty)$ is constant:

$$\frac{\mathrm{d}x_i}{\mathrm{d}t} = \kappa(\hat{x}_i - x_i) + \varepsilon_i - \frac{1}{m}\sum_{j=1}^{m} \kappa(\hat{x}_j - x_j) + \varepsilon_j \tag{S5}$$

This is the Langevin form of a stochastic differential equation in which Gaussian white noise is continuously added to deterministic dynamics (1). In particular, noise is added independently to the allocation for each item $i$. The second component of Eq. S5 enforces that capacity is constant for a given observer, by subtracting the mean change in resources. To model forgetting, when the process reaches a face of the boundary $\partial\Omega^m$ it remains confined there, this continues until vertices are reached as final absorbing states. Therefore, we have defined a truncated Ornstein-Uhlenbeck process.

The deterministic dynamics of Eq. S5 produce monotonic convergence to the desired setpoint, because the action of control is continuously scaled by the remaining distance. To illustrate, the deterministic solution for the remaining distance in each component is $|x_i(t) - \hat{x}_i| = |x_i(0) - \hat{x}_i|\mathrm{e}^{-\kappa t}$. From any starting point the distance diminishes exponentially, with larger $\kappa$ always increasing the rate of correction. In the next section, we show Eq. S5 in fact defines a feedback controller that continuously corrects deviations towards the setpoint even in the presence of noise.

We conceive of the resources determining recall at the cognitive level, making few commitments about the resources' neural implementation. That is, we follow empirically successful models of working memory that find postulating the allocation of a finite resource explains interference and memory decay as items increase, with animals eventually forgetting (2). Our model captures these broad features of memory without detailing the degree to which they are underpinned by neural spiking or changes in neural connectivity (3, 4). We simply assume that constraints emerge that limit performance. As we focus on the broad functions of working memory, we also abstract away how different memory systems interact (e.g., episodic and semantic memory), and distinctions in how their resources operate (5, 6). Understanding this interaction is key for future research examining how evolution guides the progression from performing a task to learning the task solution and retaining it in long-term storage. Future research should also examine how control strategies such as chunking enable more efficient use of available capacity (7), and how such strategies shape capacity evolution.

### 2 Resource allocation dynamics via optimal control theory

Our resource dynamics equation also arises naturally as an approximate solution to a family of optimal control problems (8). To illustrate, consider a deterministic scenario where control input $u$ directly adjusts to resource allocation $\frac{\mathrm{d}x}{\mathrm{d}t} = u$, but producing control input uses energy, suffering a penalty $\frac{1}{2}u^{\top}u$. Further, take the typical approach of producing a quadratic approximation of the reward function around the desired setpoint:

$$R(x) \approx R(\hat{x}) + \frac{1}{2}\,(\hat{x} - x)^{\top} \nabla_x^2 R(\hat{x})(\hat{x} - x) \tag{S6}$$

Continuing the illustration, let the objective be maximizing the average reward over a time period $T$:

$$\frac{1}{T}\int_0^T \left(R(\hat{x}) + \frac{1}{2}\,(\hat{x} - x)^{\top} \nabla_x^2 R(\hat{x})(\hat{x} - x) - \frac{1}{2}u^{\top}u\right) \mathrm{d}t \tag{S7}$$

This implies the Hamilton–Jacobi–Bellman equation for value function $V$:

$$0 = \frac{\partial V}{\partial t} + \max_u\{\frac{1}{2}(\hat{x} - x)^{\top} \nabla_x^2 R(\hat{x})(\hat{x} - x) - \frac{1}{2}u^{\top}u + \nabla_x V^{\top} u\} \tag{S8}$$

Here, $u$ must obey the constraint $\sum_{j=1}^{m} u_j = 0$ because capacity does not change. Together, this formulates a standard linear-quadratic regulator problem that can be solved using the Riccati equation (8, pp. 88-89). The resulting optimal control law is:

$$\hat{u} = \kappa A(\hat{x} - x) \tag{S9}$$

Here, the projection that keeps capacity constant is achieved by the matrix:

$$A = I_m - \frac{1}{m}\mathbf{1}_{\boldsymbol{m}}\mathbf{1}_{\boldsymbol{m}}^{\top} \tag{S10}$$

Substitution reveals that the optimal control law $\hat{u}$ is the deterministic version of our resource dynamics equation (Eq. S5) written in matrix form. In particular, $I_m$ is the identity matrix and $\mathbf{1}_{\boldsymbol{m}}$ is an all-ones vector. To disambiguate terms, what we call *cognitive control* $\kappa$ is the ability to change resource allocation, the *optimal control law* $\hat{u}$ is a policy for changing resource allocation. Taking a control theory lens, $\kappa$ will generally be a function of model parameters (e.g., $m, q, t$) that depend on assumptions about the specific scenario. However, to keep our model simple, we take $\kappa$ to be a fixed value that is set genetically that evolves as part of our evolutionary model.

### 3 Time course of cognitive processing events

We assume that the observer's initial resource allocation $x(0)$ is drawn from a uniform distribution over our state space $\Omega^m$ (a regular $m$-simplex), which is equivalent to $\frac{x(0)}{\eta} \sim \mathrm{Dirichlet}(\mathbf{1}_{\boldsymbol{m}})$. That is, we assume maximum uncertainty about the starting resource allocation (9, pp. 409-425). Before the cue arrives, the observer has no information about which item will be required for recall, so control moves resources towards the optimal allocation $\hat{x}_{\mathrm{precue}} = \left[\frac{\eta}{m}, \dots, \frac{\eta}{m}\right]^{\mathsf{T}}$. In the time after the cue arrives, control moves resources towards the updated optimal allocation $\hat{x}_{\mathrm{postcue}} = \hat{x}(\eta, m, q)$ (Eqs. S3 and S4). The cue carries information about the environment by indicating the likely target item. As cue reliability $q$ increases, so does cue-environment mutual information, as calculated explicitly in Eq. S27. Upon receiving the cue, the observer reallocates cognitive resources towards representing the cued item. Therefore, the information in the observer's representation of stored items depends ultimately on both cue information and the efficacy of their control in shifting resources. Although control processes attempt to maintain an optimal resource allocation distribution, eroding noise means that items are forgotten. In particular, from the first point in time that no resources are allocated to an item $i$ it is forgotten and cannot be reallocated. This means that $x_i = 0$ is an absorbing boundary. Given enough time, the system always reaches an absorbing state with all $\eta$ resources allocated to a single item, and 0 allocated to the others. So far, we have formulated a cognitive model that is rich enough to fit empirical trends in working memory tasks (see forerunners, 10–12).

### 4 Fitness function

We assume successful recall leads to benefits, with superior capacity or control incurring greater investment costs paid regardless of correct recall:

$$W(\eta,\kappa) = \bar{R}(\eta,\kappa) - c(\eta+\kappa) \tag{S11}$$

Here, $\bar{R}(\eta,\kappa)$ is the expected recall taken over randomness during cognitive processing due to: (1) initial resource allocations, (2) accrued noise, (3) variation in the time recall must be produced. For simplicity, we assume a separation of timescales between fast cognitive processing and slow evolution on traits, and that environmental stochasticity is stationary and non-fluctuating.

### 5 Finding evolutionary optimum

We developed an understanding of our evolutionary model's predictions by simulation and parameter sweeps. Our model of cognitive resource dynamics is not solvable in closed-form, and so we used Monte Carlo simulation. In particular, we can write the cognitive resource dynamic equation (Eq. S5) in Ito form (1):

$$\mathrm{d}x = \kappa A(\hat{x} - x)\,\mathrm{dt} + A\,\mathrm{d}B \tag{S12}$$

$$A = I_m - \frac{1}{m}\mathbf{1_m}\mathbf{1_m}^{\top} \tag{S13}$$

Here, $\mathrm{d}B$ is the stochastic increment and $A$ maintains the constraint that capacity is constant. For simulation, we can leverage the closed-form Ornstein-Uhlenbeck transition (13) to give the discretized form:

$$\Delta x = (1 - \mathrm{e}^{-\kappa\Delta t})A(\hat{x} - x) + \sqrt{\frac{1 - \mathrm{e}^{-2\kappa\Delta t}}{2\kappa}}A\varepsilon_t \tag{S14}$$

$$\varepsilon_t \sim \text{Normal}(0, I_m) \tag{S15}$$

To handle absorbing boundaries, we replace $A$ with the following matrix that reduces dimension and appropriately rescales when an item is forgotten:

$$A' = \text{diag}(a) - \frac{1}{\sum_{i=1}^{m} a_i} \text{diag}(a)\text{diag}(a)^\top \tag{S16}$$

$$a = [\mathbb{1}\{x_1 > 0\}, \dots, \mathbb{1}\{x_m > 0\}]^\top \tag{S17}$$

We assume the observer must produce recall at a random time drawn uniformly from period $T$, with the cue occurring at $T/2$. Producing order $10^6$ replicates allowed us to approximate expected recall $\bar{R}$ and thereby fitness $W$.

We used the covariance matrix adaptation evolutionary strategy algorithm (CMA-ES) (14) to find the values of capacity and control that maximize fitness $(\hat{\eta}, \hat{\kappa})$. This was used as a robust method to find optima, rather than because it has been studied as a way to understand biological evolution (e.g., Wright-Fisher process). However, it demonstrates optima are reachable by an evolutionary process. The CMA-ES converged from different initializations, consistent with our fitness function having a single maximum. The average relative standard error for $(\hat{\eta}, \hat{\kappa})$ solutions <0.05. Supplementary Code S2 contains simulation script. Dataset S1 contains a broad parameter sweep. Dataset S2 sweeps considering broad range of metabolic costs, and Dataset S3 sweeps that support findings about information-processing capability and environmental complexity. Dataset S4 sweeps for separate capacity and control costs. Code S1 contains plotting functions. Find these files within the repository.

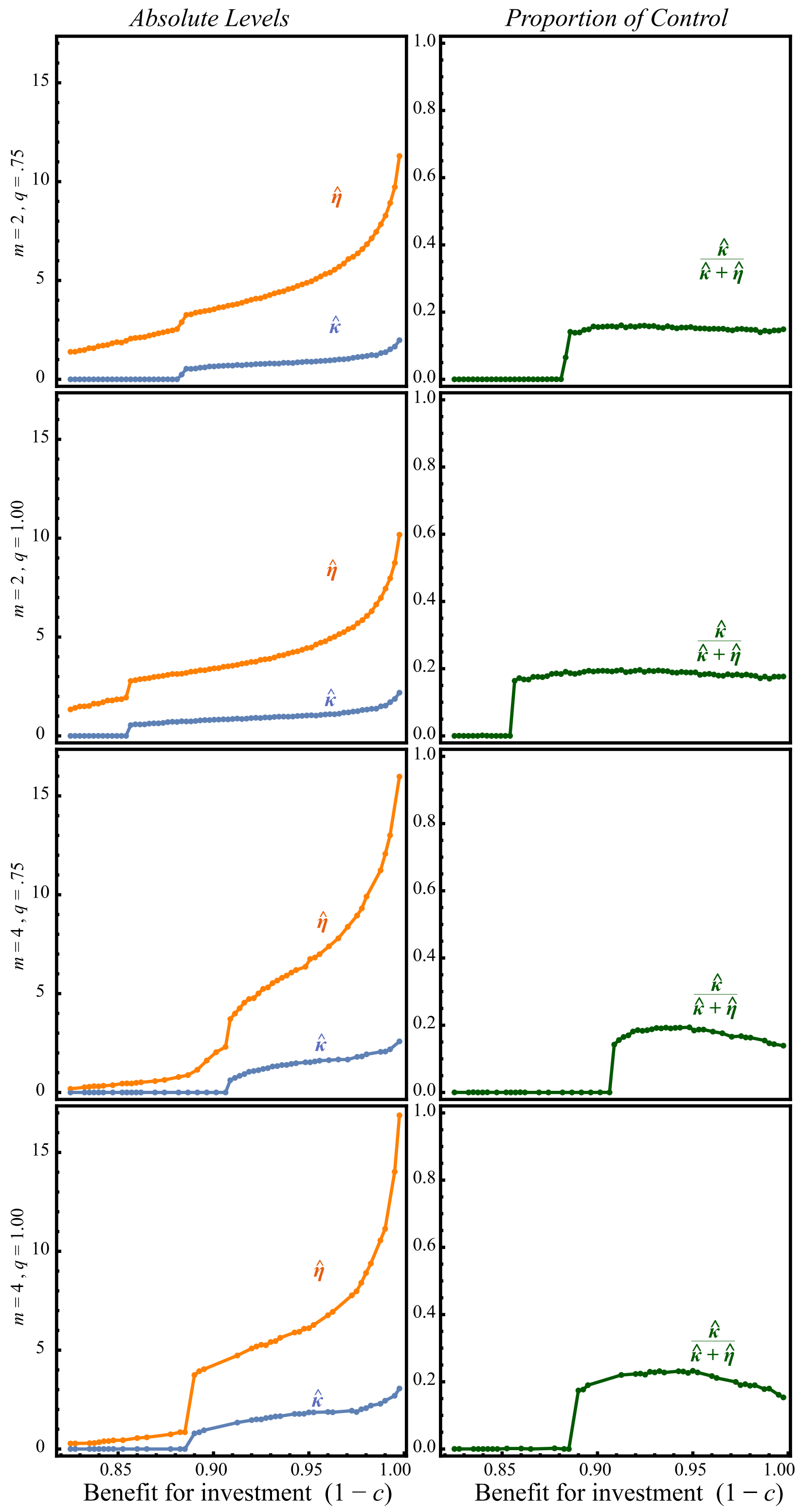


**Fig. S1.** Evolutionary capacity and control simulation optima. Examples shown for high and low values of load ($m$) and cue reliability ($q$). The average relative standard error for $(\hat{\eta}, \hat{\kappa})$ solutions <0.05, so plotted trends are highly representative. In plots, *T=3*.

### 6 Long-run expected recall approximation

We derive an approximate version of expected recall by looking at the long-run distribution of resources in the absorbing state. Only the simplest stochastic processes can be solved in closed form, and our full analysis relies on Monte Carlo simulation. However, by making several convenient assumptions we gain a closed-form solution that provides insights into how capacity and control interact. That is, the long-run expected recall $\tilde{R}$ we derive in this section can be used as a tractable approximation to the full dynamics $\bar{R}$, but is obtained under stronger assumptions. First, it considers only the 2-item case ($m = 2$), so resources are allocated to either the cued item or a non-cued item. Second, it considers the resource allocation expected in population after memories have been held for long time, and all individuals have forgotten one item and allocated all resources to the other. That is, it uses the absorbing state distribution, abstracting over transient dynamics. Nonetheless, $\tilde{R}$ provides a window into system behavior that agrees with the central conclusions from the full model.

First, we calculate the probability of ending up with resources allocated to the cued item versus the non-cued item, from a given initial starting allocation (15, pp. 664-667). Our stochastic differential equation for resource allocation dynamics (Eq. S5) can be formulated as a backward Kolmogorov equation, telling us the probability density $p$ of reaching a given state, conditional on an initial state. We change coordinate system to have a single state variable $(x_1, x_2) = (y, \eta - y)$, so that $y$ is the amount of resources allocated to the cued item. In particular, $y$ is the initial state and $y'$ state reached at a time $t$ so that $p(y, t) = \mathrm{P}(y'|y, t)$. This leads to the backward equation:

$$\frac{\partial}{\partial t}p(y,t) = -\kappa(\hat{y} - y)\frac{\partial}{\partial y}p(y,t) - \frac{1}{2}\frac{\partial^2}{\partial y^2}p(y,t) \qquad \text{(S18)}$$

Further, focusing on long-run behavior removes the time dependence and produces:

$$0 = -\kappa(\hat{y} - y)\frac{\mathrm{d}}{\mathrm{d}y}p(y) - \frac{1}{2}\frac{\mathrm{d}^2}{\mathrm{d}y^2}p(y) \qquad \text{(S19)}$$

This is a linear second-order ordinary differential equation that can be solved with suitable boundary conditions. Forgetting means that once an item is allocated no resources, it never recovers in memory. Without loss of generality, we calculate the probability mass of all resources being allocated to the cued item. This provides suitable boundary conditions: if all resources are initially allocated to the cued item, the un-cued item is instantaneously forgotten and this never changes, $p(\eta) = \mathrm{P}(\eta|\eta, t) = 1$ and $p(0) = \mathrm{P}(\eta|0, t) = 0$. Therefore, Eq. S19 can be solved by reducing the order and integrating to give:

$$p(y) = \frac{\mathrm{erfi}\left((\eta - y)\sqrt{\kappa}\right) + \mathrm{erfi}\left(\hat{y}\sqrt{\kappa}\right)}{\mathrm{erfi}\left((\eta - \hat{y})\sqrt{\kappa}\right) + \mathrm{erfi}\left(\hat{y}\sqrt{\kappa}\right)} \qquad \text{(S20)}$$

Here, erfi is the imaginary error function, a strictly increasing function of its input.

The next step is calculating the probability of all resources ending up allocated to the cued versus non-cued item, from *any* initial distribution. In the full model, the initial resource allocation is random, resource allocation dynamics unfold, and then a cue is received which further alters dynamics. The advantage of our absorbing state analysis is that it provides clear answers to questions about our model, but it must abstract over these complicated dynamics. We take the position of maximum uncertainty about how these transient dynamics unfold by assuming a uniform distribution over the support (9, pp. 409-425). That is, not knowing the true shape, we assume the maximum entropy distribution that introduces the least bias. This means the probability all resources will be allocated to the cued item is:

$$\mathrm{P}(\eta) = \int_0^{\eta} p(y) \frac{1}{\eta} dy \tag{S21}$$

$$\mathrm{P}(\eta) = \frac{\eta - \hat{y}}{\eta} + \frac{1}{\sqrt{\pi}\eta\sqrt{\kappa}} \frac{\mathrm{e}^{\hat{y}^2\kappa} - \mathrm{e}^{(\eta-\hat{y})^2\kappa}}{(\mathrm{erfi}\left((\eta - \hat{y})\sqrt{\kappa}\right) + \mathrm{erfi}(\hat{y}\sqrt{\kappa}))} \tag{S22}$$

We are now ready to assemble the long-run expected recall $\tilde{R}$. With probability $q$ the cued item must be recalled, and has all $\eta$ resources allocated to it with probability $\mathrm{P}(\eta)$, with the probability of correct recall as a function of allocated resources being $(1 - \mathrm{e}^{-\eta})$. Taking the complement of events for the non-cued item, the long run expected recall is:

$$\tilde{R} = (1 - \mathrm{e}^{-\eta})q\mathrm{P}(\eta) + (1 - \mathrm{e}^{-\eta})(1 - q)(1 - \mathrm{P}(\eta)) \tag{S23}$$

**7 Capacity and control interactions**

The approximation to expected recall from the previous Section 6, allows us to clarify more about the interaction between capacity and control. We narrow down to this approximation for model insights, which are easiest to gain if we consider a fully reliable cue ($q = 1$):

$$\tilde{R} = (1 - \mathrm{e}^{-\eta}) \frac{\mathrm{e}^{(\eta\sqrt{\kappa})^2} - 1}{\sqrt{\pi}(\eta\sqrt{\kappa})\mathrm{erfi}(\eta\sqrt{\kappa})} \tag{S24}$$

When capacity and control appear together, they do so in the term $\eta\sqrt{\kappa}$. This means that the distribution of resources $\mathrm{P}(\eta)$ is affected by capacity and control in a way that is synergistic, in the sense they scale each other. Furthermore, we see confirmation that capacity is likely to be prioritized in improving recall, because the effect of control is diminished by the square root. Additionally, capacity directly affects recall because the more resources allocated to an item the better recall $(1 - e^{-\eta})$, whereas control has only an indirect effect via altering the resource distribution.

This synergy alongside the greater efficacy of capacity is evident in Fig. S2, which extends on the main text to show the derivative of long-run expected recall with respect to capacity. As can be seen using the legend, both derivatives are positive implying synergy. The derivative with respect to capacity (i.e. the marginal returns) is larger over a greater range of values, so that greater investment in capacity is likely profitable under a broad range of cost structures. Finally, we can see that the highest returns happen at intermediate values of capacity and control, so the strength of the synergy is non-monotonic. Supplemental Dataset S2 contains parameter sweeps showing that the production of our three regimes is robust, plotting code is in Code S1.

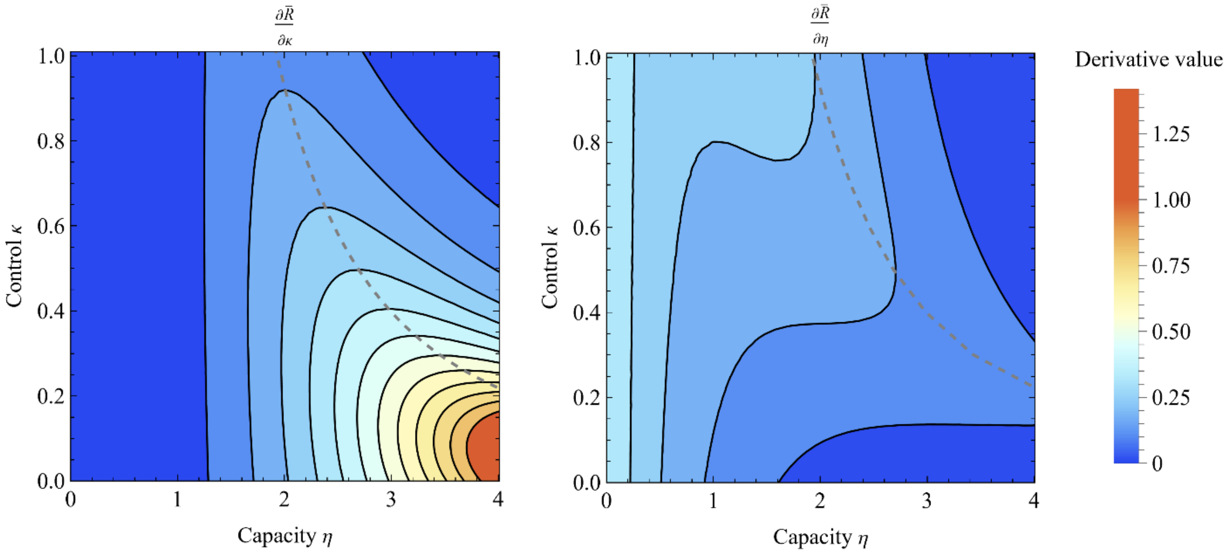


**Fig. S2.** Efficacy of recall with a change in control $\partial\tilde{R}/\partial\kappa$ and capacity $\partial\tilde{R}/\partial\eta$.

**8 Metabolic costs and why capacity is more efficacious than control**

Our evolutionary model found that capacity is broadly prioritized for investment. In the previous Section 7, we showed that a central reason for this is that increasing the reservoir of resources raises the ceiling on the potential for recall. By contrast, the ability to rearrange resources is contingent, and must make use of the reservoir that is available. A corollary is that passive storage can evolve, where capacity exists without control. Here, we explain that this capacity prioritization would arise in a wide range of stochastic processes, in which deterministic forces that attempt to revert to a point, alongside additive linear Gaussian noise (1). That is, the finding that capacity should take on larger values than control emerges fundamentally from how capacity and control are conceived of in our model: *capacity* setting the volume of total resources, with *control* being the rate of reversion to the optimal allocation against corrupting noise. Storing information entails metabolic costs due to maintaining the neural substrate, whereas reallocating information is costly because it requires modulating activity (16). Our model exemplifies a broad fact about similar stochastic systems. Whether the reverting force results from tightening a spring, or from gravity pulling a dust particle to the bottom of a water glass, there are diminishing returns such that doubling the force's strength much less than halves the amplitude of the noise.

To illustrate, consider a general Ornstein-Uhlenbeck process. There are closed-forms for the expected value and standard deviation for this process (13):

$$\mathbb{E}[x(t)|x(0)] = \hat{x}(1 - \mathrm{e}^{-\kappa t}) + x(0)\mathrm{e}^{-\kappa t} \qquad \text{(S25)}$$

$$\mathrm{SD}[x(t)|x(0)] = \sqrt{\frac{(1-\mathrm{e}^{-2\kappa t})}{2\kappa}I_m} \tag{S26}$$

In our model, $\hat{x}$ is the desired setpoint that maximizes recall, so the signal-to-noise ratio is the expected trend in resource allocation as it approaches a desired value, over the stochastic spread due to noise, $\|\mathrm{SD}[x(t)|x(0)]^{-1}\mathbb{E}[x(t)|x(0)]\|$. After substituting, we find that capacity and control affect the signal-to-noise such that $\|\mathrm{SD}[x(t)|x(0)]^{-1}\mathbb{E}[x(t)|x(0)]\| \sim \eta\sqrt{\kappa}$. The term $\eta\sqrt{\kappa}$ describes how capacity and control synergize in their effect on recall, which is why it also appears in Eq. S24. That is, capacity linearly increases signal-to-noise, whereas control improves recall by pushing the system's state against noise and therefore is suppressed by the square root. This square root relationship is a fundamental feature of systems where a linear corrective drift opposes a diffusive process (1). It arises because control acts by decreasing dispersion, which is naturally related to the squared distance around the optimum allocation. Therefore, prioritizing investment in capacity should emerge in a range of models.

Given that capacity and control synergize according to $\eta\sqrt{\kappa}$, the metabolic cost of control would have to be substantially cheaper than capacity for our prediction to be reversed with control being prioritized for investment. This is a particularly strong requirement in cases where species invest heavily in cognition. In our base model, we assumed the simplest exchange rate between a unit of investment in capacity and control, with a single linear cost parameter $c$. As expected from our mathematical analysis, this model found prioritization of investment in capacity was broadly favored. That is, much less than half of relative investment went to control, assuming equivalent per-unit costs (Fig. 3D). We further examined an extension to the model with separate linear costs, capacity $c_\eta$ and control $c_\kappa$. This robustly supported the prediction of capacity prioritization (Fig. S3). Consistent with this hypothesis, our preliminary experimental data also found both humans and rhesus monkeys displayed more inferred units of capacity than control. Nonetheless, our work has opened a path to test how broadly this holds, and the conditions for reversal that control be exceedingly cheap. Testing our model in the context of empirical measurements of metabolic costs appears an enlightening next step. For instance, developing a large brain densely populated by neurons may be more costly than the transient neural modulation and gating that may underpin attention (17).

We formally define capacity and control in terms of natural variables that arise within our dynamic systems model. Further, we assume simple linear metabolic costs, minimally capturing the notion that costs increase proportionally with investment in these natural variables. This lays the groundwork for other instantiations. For instance, defining capacity and control directly in terms of information-theoretic quantities, or considering nonlinear biologically-inspired cost functions.

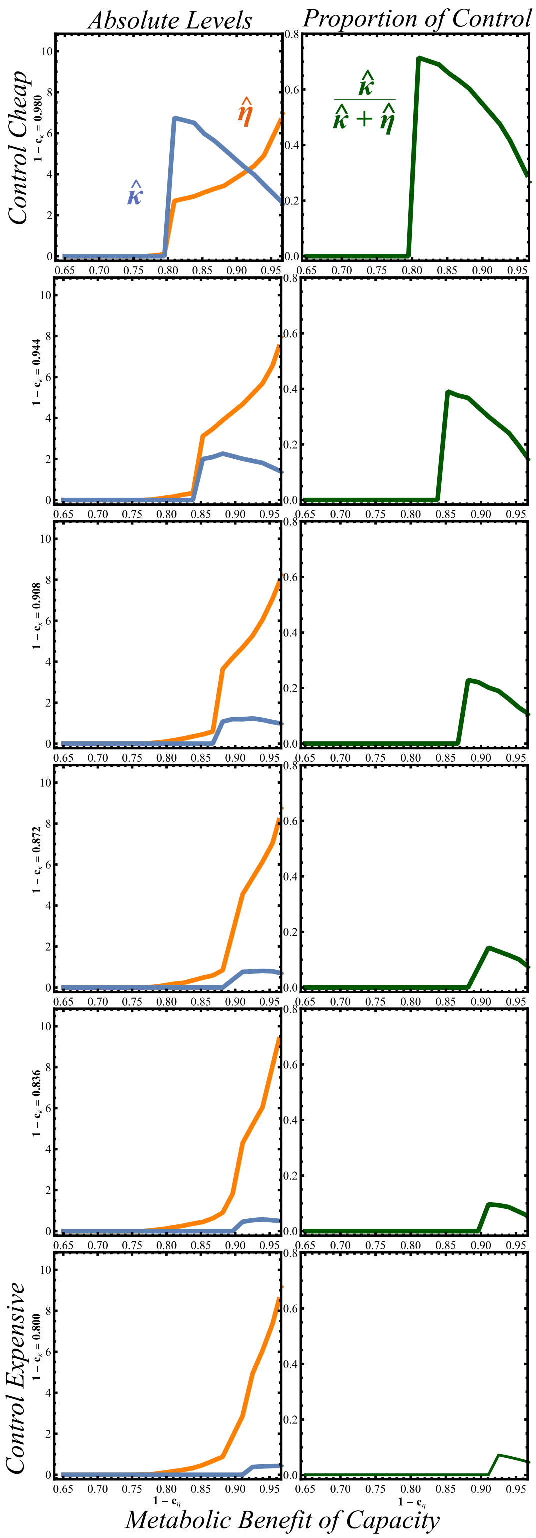


**Fig. S3.** Separate capacity and control costs. Control cost varies down rows, capacity costs vary on horizontal axis. Costs on a cognitive component reduce that component. Capacity takes on larger values than control, unless control is extremely cheap. Three regime structure emerges. Plot sets $(m, q, T) = (4,1,3)$.

### 9 Scale of capacity and control

Capacity $\eta$ is a dimensionless quantity representing the amount of cognitive resource that sets the maximum possible allocation to any item $x_i$ or the total amount across all items. Given the recall function for a given item $F(x_i) = 1 - \mathrm{e}^{-x_i}$, capacity has a concrete meaning in terms of increasing recall; the first unit of capacity allocated increases recall from 0 to $1 - \mathrm{e}^{-1}$, the second from $1 - \mathrm{e}^{-1}$ to $1 - \mathrm{e}^{-2}$, and so on. Control $\kappa$ has units of $[\mathrm{time}]^{-1}$ so is a rate constant, which is clear from our resource dynamics equation (Eq. S5). By changing the rate of approach of resources $x(t)$ to the optimal allocation, control $\kappa$ is related to observable trends by being involved in changing recall over time. As $T$ is the maximum time period, $\kappa T$ is a dimensionless quantity that gives the amount of control action that can happen within that period.

**10 Experimental test methods**

Our experimental study was preregistered prior to the collection of data (https://osf.io/gnf6h). The human study protocol was approved by the Princeton University Institutional Review Board. Behavioral data and statistical analysis code is available through repository.

**Participants.** Participants were recruited through the online platform Prolific (https://www.prolific.com/) and were required to be English-speaking adults (≥ 18 years) residing in the United States and to have ≥90 % approval ratings. Only participants using desktop or laptop computers were allowed. All participants provided informed consent online before beginning the task. Participants were compensated $5.25 USD for the 30-min experiment and received a performance-contingent bonus of $0.015 USD per correct response.

Following our preregistered sampling plan, we collected 360 participants. Of these, 14 participants were excluded based on pre-defined exclusion rules (Figs. S4 and S5). The exclusion rules aimed to remove participants that either had near-chance performance or demonstrated a high degree of bias in their response (e.g. tended to always report that they had seen the image). To identify these characteristics, we computed for each participant the signal detection theory metrics, sensitivity ($d'$) and bias (criterion, $c$). These metrics were measured by using both cue valid and invalid trials, along with non-match trials (Fig. 4), to compute a hit rate and false-alarm rate using the standard log-linear (add-0.5) correction (18, 19). We then excluded participants with either near-chance sensitivity ($d' < .25$) or high bias ($|c| > .50$). This resulted in the removal of 14 participants, leaving us with 346 for our main analysis. We included the data of all

10 rhesus macaques (*Macaca mulatta*) collected previously who took part across 6 experiments conducted by Brady & Hampton (20), who were outside our exclusion ranges.

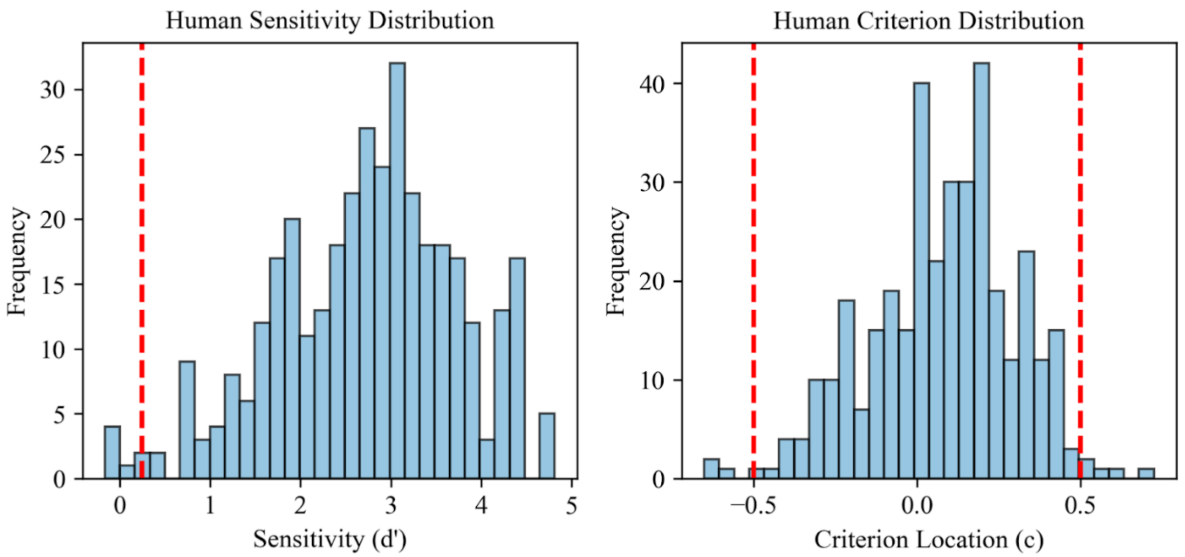


**Fig. S4.** Distribution of sensitivity and criterion values for human participants. In our main analysis, we only analyze cue valid and invalid trials (probed image was in the original set), it is therefore important to remove participants who show extreme bias in their responses, as this could mask as higher or lower overall accuracy. Additionally, we remove participants that display near chance accuracy. To separate bias from sensitivity, we use both match and non-match trials to compute signal detection theory measures of sensitivity ($d'$) and bias (criterion, $c$) for each participant. Based on our pre-registered thresholds, we removed participants with $d' < .25$ and $|c| > .5$ (vertical dashed lines in either plot). Overall, 7 human participants were removed based on $d'$ and 8 were removed based on criterion (with one participant being flagged for both), for a total of 14 removals out of 360 participants.

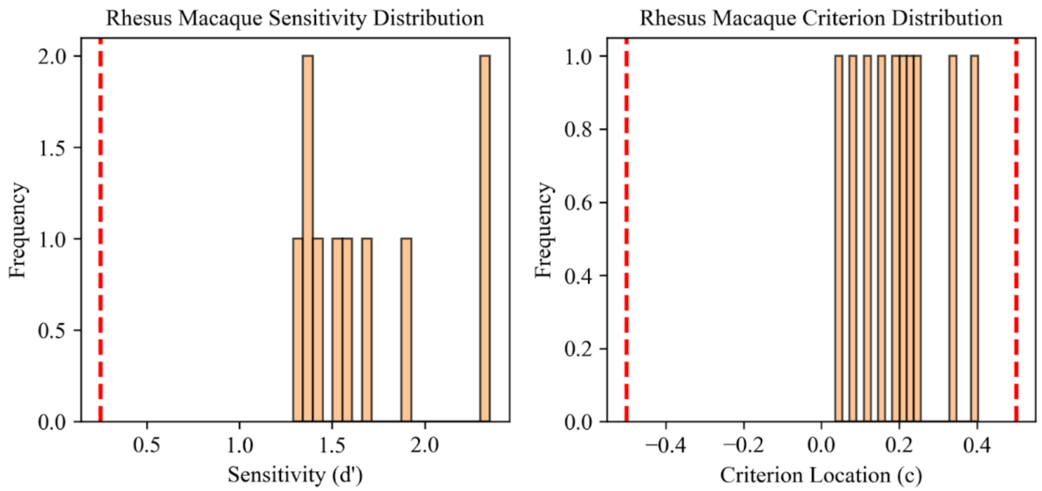


**Fig. S5.** Distribution of $d'$ and $c$ (criterion) values for rhesus macaques. Vertical lines show exclusion thresholds (none removed).

**Stimuli.** For comparability with the monkey data we used the original stimulus set provided by Brady & Hampton. The items to be recalled were drawn from nine colored photographs of natural scenes. Each image subtended 146 × 139 px and was presented within a 218 × 196 px white frame on a black background. Four peripheral frames were arranged around a central frame in a cross configuration. On each trial two, three or four images were presented simultaneously at distinct peripheral locations, depending on the load condition. The same stimuli were used for all participants, but their positions and assignments to trial types were randomized. Stimuli were presented via the *jsPsych* library (v 7.3.1) (21) running in participants' web browsers.

**Trial structure.** To ensure comprehension, participants first read written instructions followed by a short instruction quiz. Participants were required to answer every quiz question correctly before beginning the task. In the task, each trial followed the sequence of events used by Brady & Hampton (Fig. 4):

1. Ready prompt: A ready screen at the beginning of every trial, shown as a green rectangle at fixation.
2. Encoding phase: Two, three or four images were presented simultaneously at distinct peripheral locations for 400 ms.
3. Pre-cue delay: A blank screen was shown for 300, 700 or 1100 ms.
4. Retro-cue: A dashed frame blinked twice around one peripheral location. Each blink lasted 150 ms on, 150 ms off and 150 ms on (total 450 ms).
5. Post-cue delay: A second blank interval of variable duration followed. The sum of pre-cue and post-cue delay was held constant at 1,350 ms.
6. Recall test. A single test image appeared at screen center; participants had up to 3,000 ms to press "J" if the image had appeared in the study array or "F" if it was new. Correct responses triggered a "correct" tone and displayed + $0.015 for 1.5 s; incorrect responses played an error tone and displayed $0.00 for 1.5 s.
7. Inter-trial interval. After the 1.5 s feedback display, a blank interval of 3.5 s followed correct responses (total 5.0 s ITI), and a blank interval of 5.5 s followed incorrect responses (total 7.0 s timeout) before the next trial started.

Our statistical model makes predictions about accuracy on cue valid or invalid trials where the recall test image matched an image in the encoding array. Non-match trials are used to exclude participants (as outlined in *Participants*).

Our study comprised the following independent variables, which pilot testing suggested would support estimating capacity and control:

1. Cue validity: On valid trials, the cued location accurately signaled which image would be probed. On invalid trials, the cued location did not match the image that was probed. This was varied within-participant
2. Cue reliability $q$: The probability that a match trial was valid or invalid. Cue reliability was varied between-participant, being either .70 or .92, and held constant for all trials for a single participant.
3. Load $m$: The number of images the participant needed to remember, being either 2, 3 or 4 images. We used a mixed approach to manipulation to improve power while ruling out interference effects. For half of sampled participants, load was varied within-participant (in 40 trial blocks; order counterbalanced across participants). In the other half, load was varied between-participants (and held constant for a single participant).
4. Cue time (Pre-cue delay) was the amount of time between stimulus offset and the cue. This was either 300 ms, 700 ms, or 1100 ms. Cue time was manipulated in the same mixed way as load but in the opposite half of the sample. In the model, this manipulated the proportion of the total time period $T$ before the cue.
5. Species: comprised of rhesus monkeys and humans.

Note that in cases where a variable (either load or cue time) was manipulated block-wise within participant, the length of that block was always 40 trials and included 20 match and 20 non-match trials. Valid and invalid trials were distributed as evenly as possible across the three blocks, such that in $q = .70$ case, there were 6 invalid trials within each block and in the $q = .92$ case there was 1 invalid trial in the first block and 2 invalid trials in the second and third block. Block order was counterbalanced across participants.

### 11 Experimental test statistical analyses

**Visualization of performance over conditions.** To illustrate participant performance, we computed mean accuracy for each participant for each trial-type in which they participated. Fig. S6 shows across-participant mean accuracy in each trial-type with 95% between-participant confidence intervals. To highlight the effect of cue time, load and cue validity, and interactions of these with cue validity, we provide additional plots (Figs. S8, S9 and S10). For these, we computed within participant accuracy for a given cue time (or load or cue validity) for valid and invalid trials, collapsing over the other two variables, and compute between-participant mean and 95% confidence intervals over those means.

**Generalized linear mixed effects model.** Preliminary to our main statistical model, we fit separate binomial generalized linear mixed effects models of human and rhesus macaques recall performance on match trials (Tables S1 and S2). Models were fit using the MixedModels.jl package for the Julia programming language (22). For humans, we fit a model predicting trial accuracy as a function of Cue Validity, Cue Reliability, Cue Time, Load, and the interaction of Cue Validity with Cue Reliability, Cue Time and Load. Random effects were included for intercepts and for all slopes, except for those involving Cue Reliability, which was only manipulated between-participant. For rhesus macaques, because cue reliability was not varied, we fit the same model, but removed terms involving cue reliability. Model coefficients' *p*-values were obtained from the package and reflect Wald tests using a standard normal approximation.

**Statistical model estimating capacity and control.** Capacity and control parameters for the recall model were estimated for either species using hierarchical Bayesian inference, such that group-level priors for either species were used to regularize participant-level estimates for participants from that species. The joint posterior over group and participant-level parameters was approximated using No-U-Turn Sampling (23) as implemented in Stan (24). Four chains with 2000 samples (1000 discarded as burn-in) were run for a total of 4000 posterior samples. Chain convergence was assessed using Gelman-Rubin statistics and sampler diagnostics: all finite $\hat{R}$ values for sampled model parameters were below 1.01, and there were no divergent transitions. Effective sample sizes were generally adequate; among the monitored group-level parameters, the lowest bulk/tail ESS was for $\sigma_{human,\kappa}$ (~365), with other lower values for $\sigma_{human,\eta}$ (~487) and Cholesky correlation terms ($L_\rho$; ~550). A full description of the parameterization and implementation of the model and choice of priors can be found in SI Appendix, Section 13. From each drawn sample, we extracted species-level posterior distributions for (log-transformed) mean capacity and control values as well as how they vary across participants within each species. These posterior distributions were then used to calculate derived quantities and between-species contrasts (see SI Appendix, Section 13 for full model and computation of derived quantities).

**Posterior predictive accuracy plots.** We generated posterior predictive summaries of accuracy using parameter draws from the fitted hierarchical model. To do this, we drew 200 samples of species-level group mean, scale parameters and correlation factor. For each selected draw of species-level parameters, we sampled sets of participant-level parameters equal to the number of individuals in the sample. This set of species-level parameters was used to compute expected accuracy on each trial-type performed by that species. Specifically, for each condition (specified by cue reliability, cue timing and load), we computed expected accuracy for each condition for valid and invalid trials in that condition, for all sampled participants in that species and averaged across participants to obtain the species-level posterior predictive mean for that draw. We aggregate the draw-wise means to show the posterior predictive mean and the 95% credible interval over draws.

### 12 Experimental test results

In this section we report the results of our experimental test, in which we fit the cognitive model to human and rhesus macaque recall data. Tables S1 and S2 report binomial generalized linear mixed-effects models of recall accuracy for humans and rhesus macaques, respectively. Fig. S6 shows observed mean accuracy for both species across all conditions, and Fig. S7 shows the corresponding posterior predictive accuracy generated by the model. Figs. S8, S9 and S10 then compare observed and model-predicted accuracy as a function of load, pre-cue timing, and cue reliability, each in interaction with cue validity.

**Table S1.** Results from binomial generalized linear mixed effects model of human recall performance. We fit a binomial generalized linear mixed effects model to predict trial accuracy as a function of Cue Validity, Cue Reliability, Cue Time, Load, and the interaction of Cue Validity with Cue Reliability, Cue Time and Load. Random effects were included for intercepts and for all slopes, except for those involving Cue Reliability, which was only manipulated between-participant. The model predicts negative effects of Load and positive effects of cue validity on accuracy. Additionally, it predicts that cue validity interacts with cue reliability, cue time, and load. We observed significant main effects of load and cue validity, in the direction predicted by the model. With regard to interactions, we noted in our pre-registration that based on prior simulation of the model, we were likely underpowered to observe these effects. Despite this, we observed significant interactions of cue validity with cue reliability and also cue time. Furthermore, although non-significant, the other interaction effect is in the predicted direction.

| **Predictor** | **Coef.** | **SE** | **z** | **P** |
|---|---|---|---|---|
| Intercept | 2.13 | 0.06 | 33.76 | < .001 |
| Cue Validity | 1.51 | 0.10 | 15.77 | < .001 |
| Cue Reliability | −2.45 | 0.56 | −4.34 | < .001 |
| Cue Time | 0.06 | 0.12 | 0.52 | .602 |
| Load | −0.32 | 0.05 | −6.20 | < .001 |
| Cue Validity × Cue Reliability | 5.66 | 0.84 | 6.70 | < .001 |
| Cue Validity × Cue Time | −0.58 | 0.20 | −2.85 | .004 |
| Cue Validity × Load | 0.06 | 0.09 | 0.71 | .476 |

**Table S2.** Results from binomial generalized linear mixed effects model of Rhesus Macaque recall performance.

| **Predictor** | **Coef.** | **SE** | **z** | **P** |
|---|---|---|---|---|
| Intercept | 0.93 | 0.08 | 11.32 | < .001 |
| Cue Validity | 0.19 | 0.13 | 1.47 | .142 |
| Cue Time | −0.51 | 0.20 | −2.50 | .012 |
| Load | −0.12 | 0.14 | −0.89 | .371 |
| Cue Validity × Cue Time | −1.01 | 0.72 | −1.41 | .159 |
| Cue Validity × Load | −0.64 | 0.26 | −2.51 | .012 |

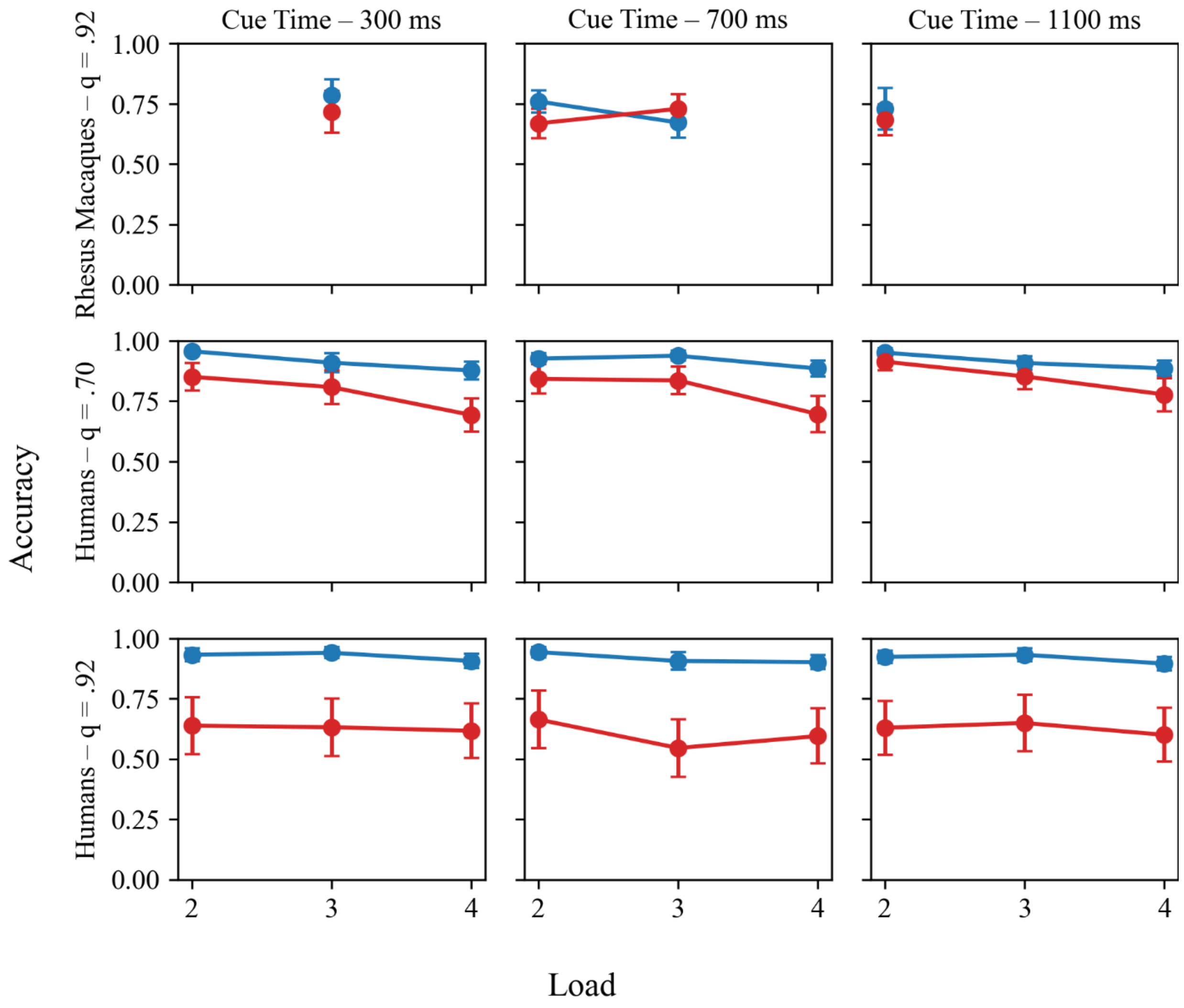


**Fig. S6.** Mean accuracy of humans and rhesus macaques on each condition. Error bars reflect 95% confidence intervals. Means and confidence intervals are computed between participants.

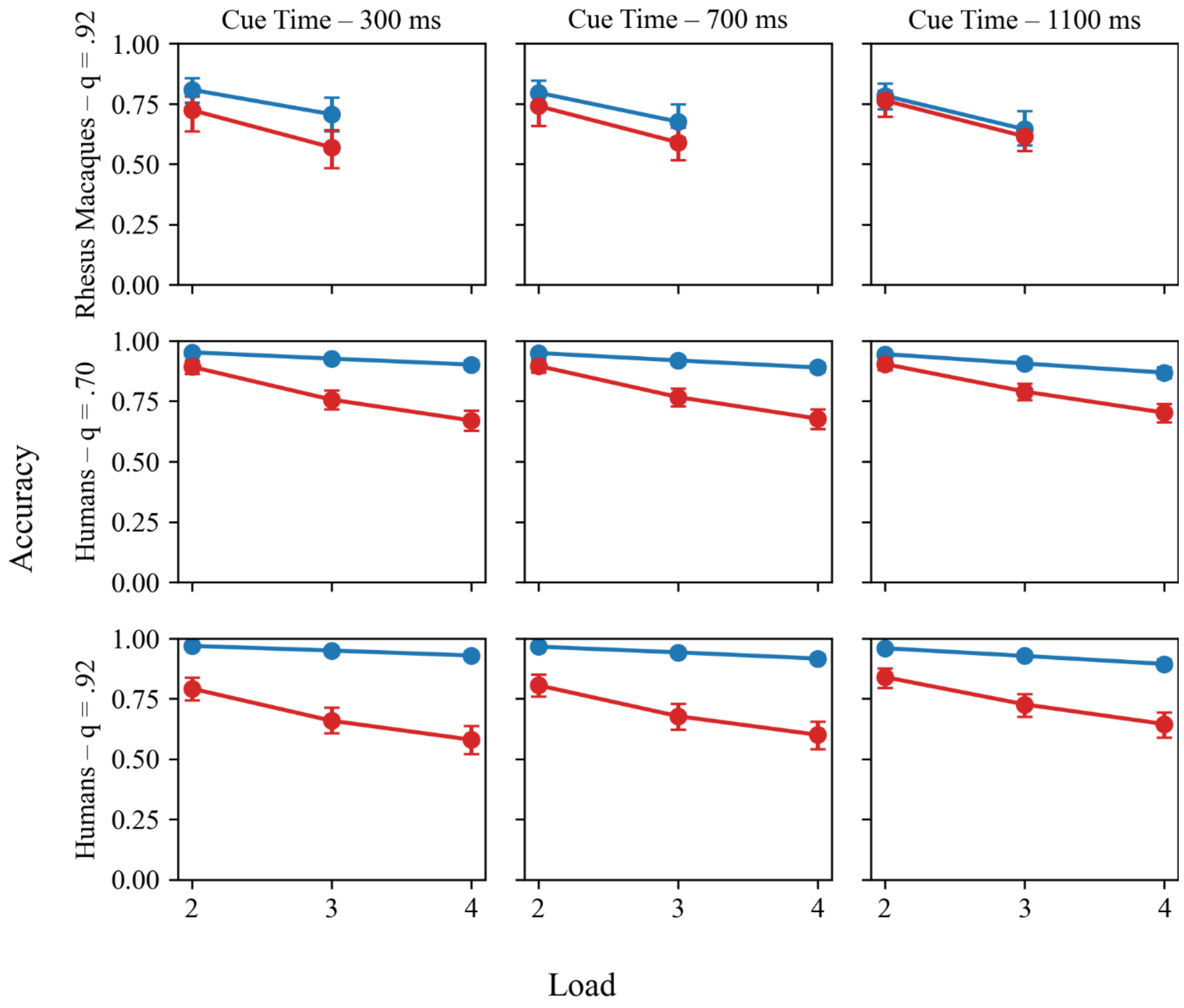


**Fig. S7.** Posterior predictive accuracy of humans and rhesus macaques on each condition, from model fit to data.

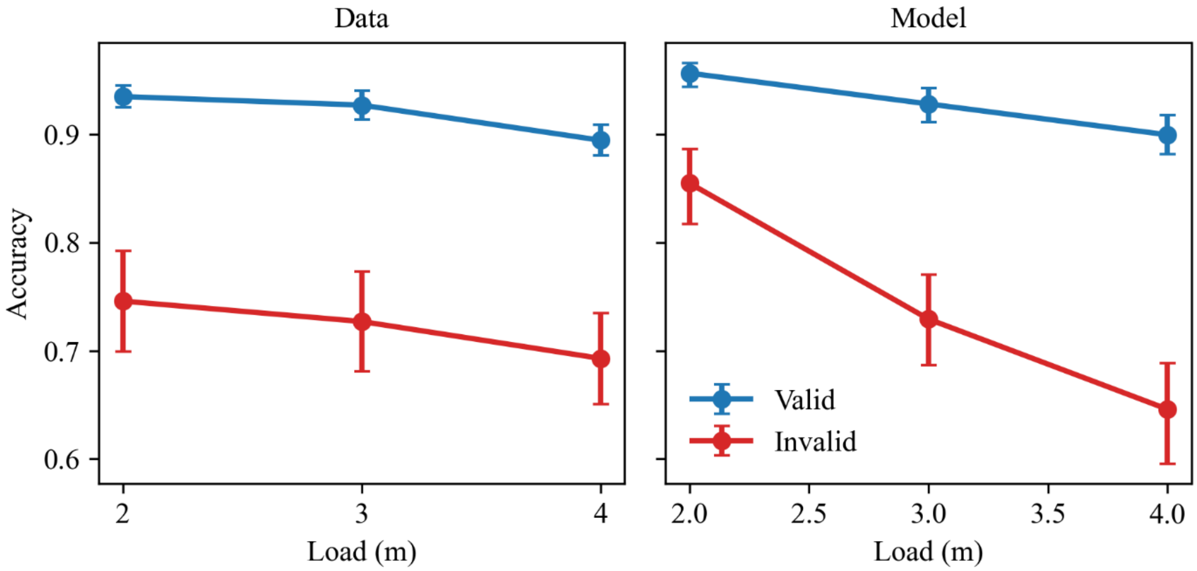


**Fig. S8.** Mean accuracy on match trials as a function of load and cue validity for Human Participants (left) and Model posterior predictions (Right). Mean accuracy collapses over cue reliability and cue timing. The model predicts both a negative main effect of load as well as an interaction of load and cue validity. We detected a main effect of load (Table S1). The interaction effect of load and cue validity was in the predicted direction but was not statistically significant.

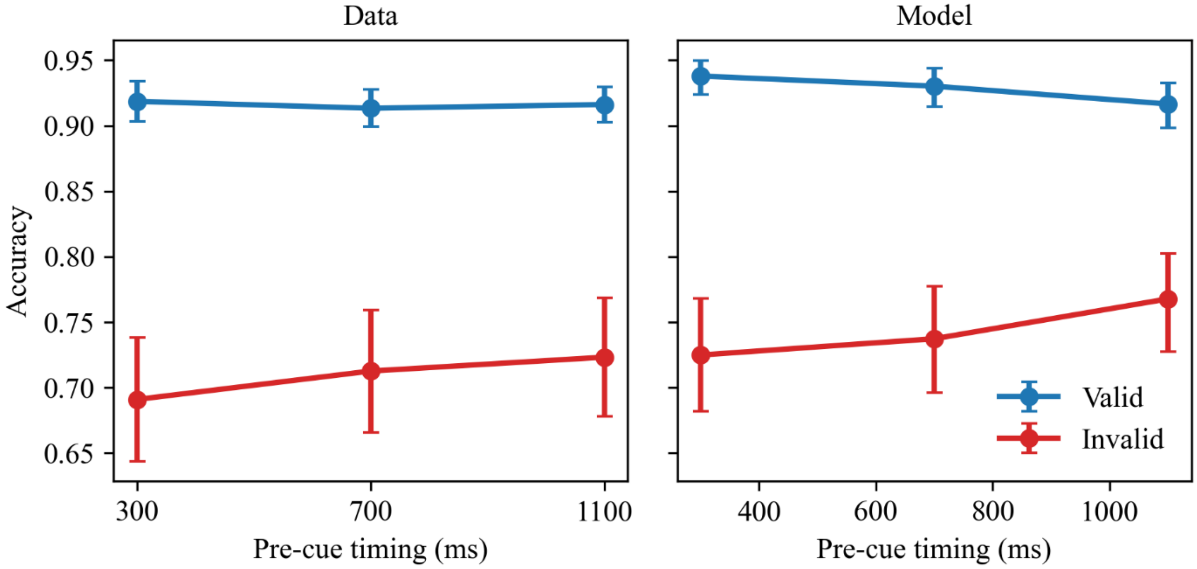


**Fig. S9.** Mean accuracy against pre-cue timing and cue validity for humans (left) and model posterior predictions (Right). Mean accuracy collapses over cue reliability and load. The model predicts a negative interaction of cue validity and pre-cue timing. We detected a main effect of cue validity (Table S1) and a significant interaction of cue validity and cue timing.

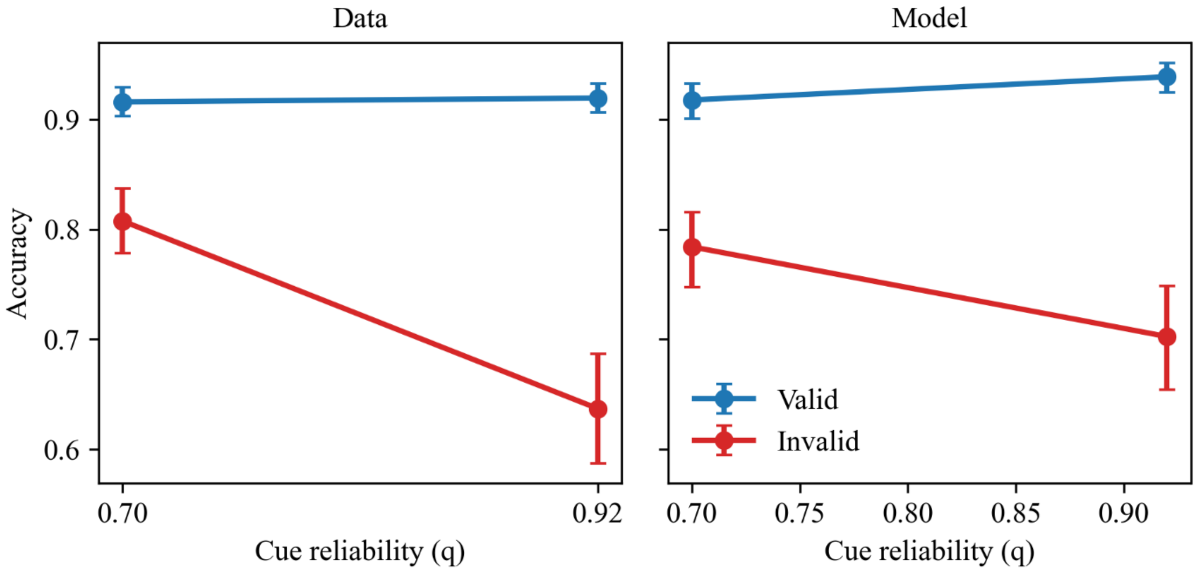


**Fig. S10.** Mean accuracy on match trials as a function of cue reliability and cue validity for human participants (left) and model posterior predictions (Right). Mean accuracy collapses over cue timing and load. The model predicts a positive interaction of cue reliability and cue validity. We observed a significant interaction of cue reliability and cue validity in human data (Table S1).

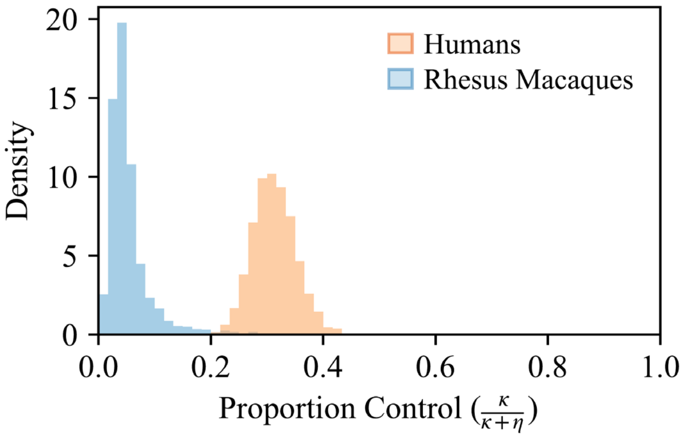


**Fig. S11. Proportion investment in control for either species.** Humans invest more in control as a proportion of their total investment than rhesus macaques. Figures show posterior distributions over the proportion of total investment that members of either species invest in control. For both humans and rhesus macaques, at the group-level, capacity exceeded control (Posterior Capacity - Control in Humans: M = 6.07; 95% CrI [4.51, 7.70]; Rhesus Macaques: M = 4.16; 95% CrI [3.25, 5.26]). Humans however, invested more in control as a proportion of total investment than Rhesus Macaques (Posterior Difference in Proportion Control, Humans - Rhesus Macaques: M = .26; 95% CrI [0.13, 0.35]).

### 13 Full statistical model and derived quantities

To estimate the capacity and control of humans and monkeys, we fit the recall model to retro-cue performance data. Our statistical model tested predictions of the theoretical model by implementing the expected recall function $\overline{R}$ (SI Appendix, Sections 1-3). Because direct Monte Carlo marginalization is computationally intensive and incompatible with deterministic likelihood optimization, we use a neural-network surrogate of $\overline{R}$ (25). The statistical model was fit with Stan (24); the neural network was trained in Scikit-learn (26).

**Expected recall.** We take the expected recall of a participant, $j$, to be a function of their capacity and control, as well as condition:

$$\overline{R}_j\colon\ \left(\eta_j, \kappa_j, \nu;\ q, t_{\text{pre}}, m, k\right) \to [0,1).$$

$\overline{R}_j$ is the expected recall accuracy of the process described in SI Sections 1-3. Here $k$ denotes cue validity; $q$ denotes cue reliability, $t_{\text{pre}}$ is pre-cue time and $m$ denotes load, $\eta_j$ denotes capacity of participant $j$, $\kappa_j$ denotes control of participant $j$. To adapt the model to fully describe human and monkey experimental data, we introduce an additional parameter, $\nu$, which controls per-timestep noise in the diffusion process, by scaling each noise term in (S5) by $\sqrt{\nu}$. To improve identifiability of $\eta_j$ and $\kappa_j$, we assume $\nu$ does not vary across participants and is sampled from the following prior distribution,

$$\log(\nu) \sim \mathcal{N}(.55,\ 0.75^2).$$

This specific prior was chosen so that it would be centered at the geometric mean of the surrogate bounds for $\nu$ (described below), so as to keep all prior mass on $\nu$ around where the surrogate was trained, and to provide a spread that would be weakly informative.

**Surrogate for $\overline{R}$ (likelihood approximation network).** There is no simple closed form for $\overline{R}_j$ because it requires marginalizing over noisy events at each time-step (Eq. S5). We therefore approximate $\overline{R}_j$ with a simulation-trained surrogate (25). To train the surrogate, we first simulate a large dataset by sampling 30,000 parameter triples $(\eta, \kappa, \nu)$. For each triple and for each task condition $(q, t_{\text{pre}}, m, k)$, we computed $\overline{R}$ as the mean of 800 noisy simulations. We then trained a feedforward multi-layer perceptron (two hidden layers, 128 units each) to predict $\overline{R}$ from $(\eta, \kappa, \nu, q, t_{\text{pre}}, m, k)$, minimizing mean squared error on logit-transformed probabilities. The resulting network provides a deterministic, computationally efficient surrogate that we embed in the likelihood.

**Observation model**. To separate attentional lapses from extra-binomial variability, we use two components. First, we mix the surrogate-predicted accuracy with chance:

$$\mu_{j,k,q,t_{\text{pre}},m} = (1 - \epsilon_j)\,\overline{R}_j(k, q, t_{\text{pre}}, m) \; + \; \epsilon_j \cdot 0.5.$$

Second, we model counts with a Beta–Binomial in mean–concentration form:

$$y_{j,k,q,t_{\text{pre}},m} \;\sim\; \text{BetaBinomial}(n_{j,k,q,t_{\text{pre}},m},\; \mu_{j,k,q,t_{\text{pre}},m},\; \phi_j),$$

where $\phi_j$ is a participant-level concentration parameter (larger $\phi_j$ approaches Binomial).

Hierarchical structure for capacity and control. Capacity and control vary by species, $s$. Because $\eta > 0$ and $\kappa \geq 0$, we model them on the log scale:

$$\begin{bmatrix} \log(\eta_j) \\ \log(\kappa_j) \end{bmatrix} \sim \mathcal{N}\big(\,\boldsymbol{\mu} + \mathbf{1}[s(j) = \text{human}]\,\boldsymbol{\beta},\ \boldsymbol{\Sigma}_{s(j)}\big)$$

Here, $s(j)$ is the species of participant $j$, $\boldsymbol{\mu} = [\mu_\eta, \mu_\kappa]^\top$ is the monkey baseline; $\boldsymbol{\beta} = [\beta_\eta, \beta_\kappa]^\top$ is the human–monkey adjustment (species contrast). Priors are

$$\boldsymbol{\mu} \sim \mathcal{N}\big(\bar{\boldsymbol{\mu}},\ \sigma_\mu^2 \mathbf{I}\big), \qquad \boldsymbol{\beta} \sim \mathcal{N}\big(\mathbf{0},\ \sigma_\beta^2 \mathbf{I}\big),$$

with $\big(\bar{\boldsymbol{\mu}}, \sigma_\mu, \sigma_\beta\big) = (0,\ 2.0,\ 1.0)$. This type of parameterization has been previously recommended as a way to test for group differences (27).

Random effects enter via species-specific covariance:

$$\boldsymbol{\Sigma} = \text{diag}(\boldsymbol{\sigma_s})\ (LL^\top)\ \text{diag}(\boldsymbol{\sigma_s}),$$

where $\boldsymbol{\sigma_s} = [\sigma_{s,\eta}, \sigma_{s,\kappa}]^\top \sim \text{Exponential}(\bar{\tau})$ and $L$ is the Cholesky factor of a correlation matrix with $L \sim \text{LKJ}(\bar{L})$. We use $(\bar{\tau}, \bar{L}) = (1,\ 2)$.

**Hierarchy for dispersion and lapses**. For extra-binomial dispersion, participant-level concentration parameters have species-level priors:

$$\log(\phi_j) \sim \mathcal{N}(\bar{\theta}_{s(j)},\ \sigma_{\theta,s(j)}^2), \qquad \bar{\theta}_s \sim \mathcal{N}(\log(100),\ 1.0^2), \quad \sigma_{\theta,s} \sim \text{Exponential}(1.0).$$

These priors are chosen to put $\phi_j$ near 100, which places variance as close to Binomial, but not forced. Participant deviations are $z_{\theta,j} \sim \mathcal{N}(0,1)$ with $\log\phi_j = \bar{\theta}_{s(j)} + \sigma_{\theta,s(j)}\, z_{\theta,j}$.

Participant-level lapses follow $\text{logit}\big(\epsilon_j\big) \sim \mathcal{N}(\text{logit}(0.02),\ 1.0^2)$.

**Derived quantities and comparison.** For hypotheses, because exponentiation induces skewness in distributions over original-scale values, we focus on species-level geometric means (medians) of capacity and control on the original scale:

$$\begin{bmatrix}\bar{\eta}_s \\ \bar{\kappa}_s\end{bmatrix} = \exp(\boldsymbol{\mu} + \mathbf{1}[s = \text{human}]\ \boldsymbol{\beta}).$$

This focuses values on the typical member of a species. Fig. 5 shows 95% HDI values of $\begin{bmatrix}\bar{\eta}_s \\ \bar{\kappa}_s\end{bmatrix}$ for either species.

We additionally compute two derived species-level quantities from posterior draws. First, we define the proportion of control as the expected fraction of control relative to total resource for a randomly drawn participant of species $s$, $\mathrm{p}_\mathrm{s}^\mathrm{ctrl} = \mathbf{E}_\mathrm{j}\left[\frac{\kappa_\mathrm{js}}{(\kappa_\mathrm{js}+\eta_\mathrm{js})}\right]$. We approximate this expectation by Monte Carlo averaging across simulated participant effects for each posterior draw: $\mathrm{p}_\mathrm{s}^\mathrm{ctrl} \approx \frac{\sum_{\mathrm{j}=1}^{\mathrm{N}}\frac{\kappa_\mathrm{js}}{(\kappa_\mathrm{js}+\eta_\mathrm{js})}}{\mathrm{N}}$. Posterior distributions of $\mathrm{p}_\mathrm{s}^\mathrm{ctrl}$ for either species $s$ are shown in Fig. S11.

Second, we define information-processing capability (IPC) at a reference condition ($q^* = .92, t_{pre}{}^* = 700\text{ ms}, m^* = 3$; see Section 15, below). Expected recall at this condition is obtained from the surrogate as a mixture of valid and invalid cue probabilities averaged across simulated participants, $s$: $\bar{\mathrm{R}}_\mathrm{s}\left(\mathrm{q}^*, t_{pre}{}^*, \mathrm{m}^*\right) = \mathrm{q}^*\ \mathrm{p}_\mathrm{s}^\mathrm{valid} + (1 - \mathrm{q}^*)\mathrm{p}_\mathrm{s}^\mathrm{invalid}$.

Here, $\mathrm{p}_\mathrm{s}^\mathrm{valid}$ and invalid are the output of the surrogate on valid and invalid trial-types, given reference condition settings and parameters of participant $s$. Information-processing capability multiplies this expected recall by the available information under set size m (measured in bits):

$$\mathrm{IPC_s} = H(m^*)\,\bar{\mathrm{R}}_\mathrm{s}(\mathrm{q}^*, t_{pre}{}^*, \mathrm{m}^*)$$

$$H(m) = \log_2 m$$

Posterior distributions over $\mathrm{IPC_s}$ are shown in Fig. S13.

### 14 Model comparison of species differences in capacity and control

To provide an additional test that both capacity and control distributions differ between species, we compared a set of nested hierarchical models which varied in whether the species-level distributions of capacity $\eta$ and/or control $\kappa$ were allowed to differ between rhesus macaques and humans. Specifically, we fit the following four (nested) variants of the model that differ only in whether the group-level distribution of $\log(\eta)$ and $\log(\kappa)$ is shared across species or allowed to differ:

1. $M_{both-shared}$: both $log(\eta)$ and $log(\kappa)$ are drawn from group-level distributions that are shared between species
2. $M_{\eta\ varies}$: $\log(\eta)$ is drawn from separate species-specific distribution; $\log(\kappa)$ is drawn from a distribution shared between species
3. $M_{\kappa\ varies}$: $\log(\kappa)$ is drawn from separate species-specific distribution; $\log(\eta)$ is drawn from a distribution shared between species
4. $M_{both-vary}$: both $log(\eta)$ and $log(\kappa)$ are drawn from species-specific group-level distributions.

All other model components were identical across variants. We evaluated each variant's performance using participant-level 5-fold cross validation, holding out entire subjects at a time

(28). Following subject removal (Figs. S4 and S5), our dataset comprised 10 monkeys and 346 humans. We constructed five folds stratified by species such that each fold held 2 monkeys and 69-70 humans, with no overlap between folds and each participant held out exactly once. For each fold and each model variant, we refit the model using only non-heldout subjects and computed the held-out log predictive density for each held-out subject by summing the log likelihood contributions across that subject's trials under the posterior draws from the training fit. Specifically, for held-out subject $s$, we scored predictive accuracy by summing trialwise log predictive densities and averaging over posterior draws: $\mathrm{elpd}_s = \log\left[\frac{1}{D}\sum_{d=1}^{D}\exp\left\{\sum_{i\in s}\log p\left(y_i \mid \theta^{(d)}\right)\right\}\right]$ where $i$ indexes trials belonging to subject $s$, $y_i$ is the observed trial outcome, and $\theta^{(d)}$ is posterior draw $d$ from the fit to non-held-out subjects. The overall $k$-fold expected log predictive density was then computed as $ELPD = \sum_{s=1}^{S} elpd_s$, where each subject's $elpd_s$ comes from the fold in which that subject was held out. Because held-out random-effect draws in the fully shared model occasionally produced degenerate overdispersion values, we applied the same numerical guard to all model variants during held-out scoring, excluding posterior draws with held-out $\tau > 10^6$ before aggregating predictive densities. For paired comparison between models, we computed per-subject differences in $elpd_s$ and the total difference as the sum of these across subjects. We also report the estimated standard error (SE) of the total difference (28).

Model comparison results are reported in Table S3. We found that the variant allowing both capacity and control to vary across species $M_{both-vary}$, achieved the best out-of-sample predictive performance (largest $ELPD$). Relative to $M_{both-vary}$, variants allowing only one component to vary showed smaller $ELPD$, with modest uncertainty in the corresponding $\Delta ELPD$.

The variant with both components shared, $M_{both-shared}$, was the worst performing. Altogether, this indicates that both capacity and control contribute to species differences in recall performance.

**Table S3.** Results from model-comparison of species-differences in capacity and control.

| **Model** | $ELPD$ | $\Delta ELPD$ (relative to best model) | SE($\Delta ELPD$) |
|---|---|---|---|
| $M_{both-vary}$ | −3066.10 | 0.00 | 0.00 |
| $M_{\kappa\ varies}$ | −3071.36 | 5.26 | 2.80 |
| $M_{\eta\ varies}$ | −3073.86 | 7.76 | 3.28 |
| $M_{both-shared}$ | −3089.31 | 23.21 | 7.94 |

### 15 Complex tasks select for information-processing capability

Previous work has emphasized the importance of changes in information processing capability for explaining intelligence (29). Our model distinguishes pure storage capacity from the control strategies that can be used to manipulate stored representations. However, we can also explore how these two factors combine to determine overall information processing capability. We find consistency with prior evolutionary models that show complex tasks select for superior information processing (30–42).

We measure the *complexity* of a task by the number of bits required to encode it without loss $H(m) = \log_2(m)$. Therefore, we take an information-theoretic definition of complexity as the amount of uncertainty that must be resolved for perfect performance (43). Complex tasks are those for which success is unlikely in the absence of knowledge, leading to a coherent measure that can be applied across tasks faced by animals (33–35). In our model, more information is required when there are more items to remember. This maps to ecological challenges such as tracking the location of multiple food items or group members.

We measure *information-processing capability* as the overall proficiency of the observer considering the combined operation of capacity alongside control by $\bar{R}(\eta, \kappa)H(m)$. That is, often being successful at a complex task implies superior information processing. We think of capacity and control as combining to give an animal's overall information-processing capability. That is, we provide a proxy for information the observer processes about a task as expressed in correct or incorrect recall. Intuitively, the fraction of task information processed is the fraction of trials correct.

We find that solely increasing the complexity of tasks leads to rudimentary information processing, in the absence of cues that allow the tracking of task features (Fig. S12). This is primarily because when an animal must store many items, their a priori probability of correctly recalling is low, so investing in information processing generates little benefit. It is not simply the case that a challenging world selects for a larger brain.

We find informative cues are required to make rich representations profitable (Fig. S12). The mutual information between the cue and environment is:

$$q \log_2(mq) + (1-q)\log_2\left(m\frac{1-q}{m-1}\right) \qquad \text{(S27)}$$

Here, the probability a cue indicates the picked item is $q$, and indicates a non-cued item with probability $\frac{1-q}{m-1}$. This is the mutual information for an $m$-ary symmetric channel with uniform input and accuracy $q$ (9, p. 190). When an observer receives a cue about which item is likely to lead to a reward, greater control is selected because it now pays to reallocate resources. For instance, the rustle of a group member cueing that a food item is taken leads the observer to update their representation. If the scene the observer attempts to encode is also complex, there are the conditions for substantial joint investment in control alongside capacity. This is because, while the observer must grapple with a complex task that requires significant information to represent, reliable cues allow it to deploy control to gain rewards. Consequently, our model suggests the hypothesis that the amount of information provided by cues about valuable tasks corresponds to the animal's ability to process information.

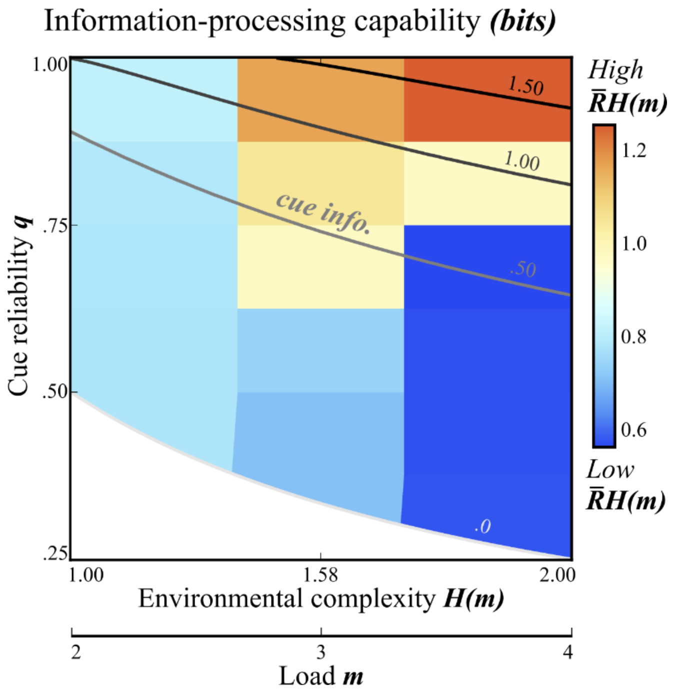


**Fig. S12.** Task complexity and cue reliability shape information-processing capability (example sets $c = .10, T = 3$). The information complexity of the task presented by the environment $H(m)$ increases with the load of m items, where $H(m) = \log_2(m)$ is the entropy of uninformed guessing. Cue reliability $q$ (vertical axis) along with the complexity of the environment (horizontal axis) determines the information carried by the cue (cross-cutting greyscale contours). Cues carry higher information in uncertain environments with reliable cues. Plotted in color are features of the animal's cognition: warmer colors indicate higher values of information-processing capability. High information-processing capability $\bar{R}(\eta, \kappa)H(m)$ evolves when there are reliable cues to a complex environment, as success often occurs, although a complex representation is required. Highly informative cues lead to higher information-processing capability. When cues are unreliable, a complex environment leads to low information-processing capability because success is rare.

We can also apply the same ideas to the analysis of our empirical data. Using the same metric for information-processing capability, combining accuracy with task complexity, we find that humans have a greater information-processing capability than rhesus monkeys (Fig. S13). Humans express nearly all 1.58 bits possible in the load 3 task. Because task complexity bounds the information processing an animal can express, difficult tasks are required to find the upper limit of a species.

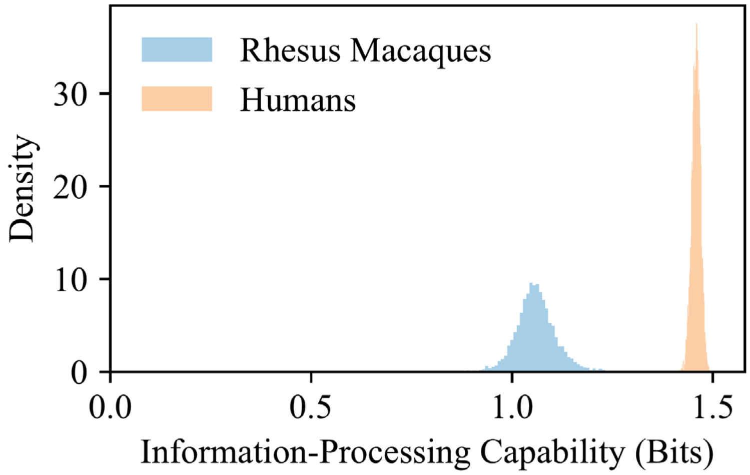


**Fig. S13**. Information-processing capability for humans and rhesus macaques. To obtain a comparable measure for humans and monkeys, we computed expected performance in the condition cue reliability .92, load 3, and pre-cue delay 700 ms, leading to $\bar{R}(\eta, \kappa)H(3)$. Humans had greater information-processing capability than monkeys (M = .40, 95% CrI [0.29, 0.50]).

**References**


1. C. W. Gardiner, *Handbook of stochastic methods*, 3rd Ed. (Springer, 2004).

2. P. M. Bays, S. Schneegans, W. J. Ma, T. F. Brady, Representation and computation in visual working memory. *Nat. Hum. Behav.* **8**, 1016–1034 (2024).

3. M. G. Stokes, "Activity-silent" working memory in prefrontal cortex: a dynamic coding framework. *Trends Cogn. Sci.* **19**, 394–405 (2015).

4. S. Schneegans, R. Taylor, P. M. Bays, Stochastic sampling provides a unifying account of visual working memory limits. *Proc. Natl. Acad. Sci. U. S. A.* **117**, 20959–20968 (2020).

5. D. Norris, Short-term memory and long-term memory are still different. *Psychol. Bull.* **143**, 992–1009 (2017).

6. B. E. Sherman, N. B. Turk-Browne, E. V. Goldfarb, Multiple memory subsystems: Reconsidering memory in the mind and brain. *Perspect. Psychol. Sci.* **19**, 103–125 (2024).

7. T. F. Brady, T. Konkle, G. A. Alvarez, Compression in visual working memory: using statistical regularities to form more efficient memory representations. *J. Exp. Psychol. Gen.* **138**, 487–502 (2009).

8. W. H. Fleming, R. W. Rishel, *Deterministic and stochastic optimal control* (Springer, 1975).

9. T. M. Cover, J. A. Thomas, *Elements of Information Theory*, 2nd Ed. (John Wiley & Sons, 2012).

10. E. Russek, *et al.*, Modeling the contributions of capacity and control to working memory development. *Proceedings of the Annual Meeting of the Cognitive Science Society* **46** (2024).

11. J. W. Suchow, D. D. Bourgin, T. L. Griffiths, Evolution in mind: Evolutionary dynamics, cognitive processes, and Bayesian inference. *Trends Cogn. Sci.* **21**, 522–530 (2017).

12. R. van den Berg, W. J. Ma, A resource-rational theory of set size effects in human visual working memory. *eLife* **7**, e34963 (2018).

13. D. T. Gillespie, Exact numerical simulation of the Ornstein-Uhlenbeck process and its integral. *Phys. Rev. E Stat. Phys. Plasmas Fluids Relat. Interdiscip. Topics* **54**, 2084–2091 (1996).

14. J. Brea, *CMAEvolutionStrategy.jl* (2023).

15. S. P. Otto, T. Day, *A Biologist's Guide to Mathematical Modeling in Ecology and Evolution* (Princeton University Press, 2007).

16. P. Sterling, S. Laughlin, *Principles of Neural Design* (MIT Press, 2015).

17. K. Kondrakiewicz, J. Nawrocka, Brains are expensive, but cognition is often cheap. *Neurosci. Biobehav. Rev.* **179**, 106450 (2025).

18. M. J. Hautus, Corrections for extreme proportions and their biasing effects on estimated values ofd′. *Behav. Res. Methods Instrum. Comput.* **27**, 46–51 (1995).

19. H. Stanislaw, N. Todorov, Calculation of signal detection theory measures. *Behav. Res. Methods Instrum. Comput.* **31**, 137–149 (1999).

20. R. J. Brady, R. R. Hampton, Post-encoding control of working memory enhances processing of relevant information in rhesus monkeys (*Macaca mulatta*). *Cognition* **175**, 26–35 (2018).

21. J. R. de Leeuw, jsPsych: a JavaScript library for creating behavioral experiments in a Web browser. *Behav. Res. Methods* **47**, 1–12 (2015).

22. D. Bates, *et al.*, *JuliaStats/MixedModels.jl: v4.25.1* (Zenodo, 2024).

23. M. D. Hoffman, A. Gelman, The no-U-Turn Sampler: Adaptively setting path lengths in Hamiltonian Monte Carlo. *Journal of Machine Learning Research* ***15***, 1593–1623 (2014).

24. Stan Development Team, "Stan Reference Manual" (2024).

25. A. Fengler, L. N. Govindarajan, T. Chen, M. J. Frank, Likelihood approximation networks (LANs) for fast inference of simulation models in cognitive neuroscience. *eLife* **10**, e65074 (2021).

26. F. Pedregosa, G. Varoquaux, A. Gramfort, V. Michel, B. Thirion, O. Grisel, … É. Duchesnay, Scikit-learn: Machine learning in Python. *J. Mach. Learn. Res.* **12**, 2825–2830 (2011).

27. M. Moutoussis, A. K. Hopkins, R. J. Dolan, Hypotheses about the relationship of cognition with psychopathology should be tested by embedding them into empirical priors. *Front. Psychol.* **9**, 2504 (2018).

28. A. Vehtari, A. Gelman, J. Gabry, Practical Bayesian model evaluation using leave-one-out cross-validation and WAIC. *Stat. Comput.* **27**, 1413–1432 (2017).

29. J. F. Cantlon, S. T. Piantadosi, Uniquely human intelligence arose from expanded information capacity. *Nat. Rev. Psychol.* **3**, 275–293 (2024).

30. P. Godfrey-Smith, *Complexity and the Function of Mind in Nature* (Cambridge University Press, 1996).

31. A. S. Dunlap, D. W. Stephens, Reliability, uncertainty, and costs in the evolution of animal learning. *Current Opinion in Behavioral Sciences* **12**, 73–79 (2016).

32. K. Aoki, M. W. Feldman, Evolution of learning strategies in temporally and spatially variable environments: a review of theory. *Theor. Popul. Biol.* **91**, 3–19 (2014).

33. M. C. Donaldson-Matasci, C. T. Bergstrom, M. Lachmann, The fitness value of information. *Oikos* **119**, 219–230 (2010).

34. R. K. Pike, J. M. McNamara, A. I. Houston, A general expression for the reproductive value of information. *Behav. Ecol.* **27**, 1296–1303 (2016).

35. O. Rivoire, S. Leibler, The value of information for populations in varying environments. *J. Stat. Phys.* **142**, 1124–1166 (2011).

36. M. Ammar, L. Fogarty, A. Kandler, Social learning and memory. *Proc. Natl. Acad. Sci. U. S. A.* **120**, e2310033120 (2023).

37. S. Dridi, L. Lehmann, Environmental complexity favors the evolution of learning. *Behav. Ecol.* **27**, 842–850 (2016).

38. B. Kerr, M. W. Feldman, Carving the cognitive niche: optimal learning strategies in homogeneous and heterogeneous environments. *J. Theor. Biol.* **220**, 169–188 (2003).

39. T. J. H. Morgan, J. W. Suchow, T. L. Griffiths, Experimental evolutionary simulations of learning, memory and life history. *Philos. Trans. R. Soc. Lond. B Biol. Sci.* **375**, 20190504 (2020).

40. K. N. Laland, J. R. Kendal, "What the models say about social learning" in *The Biology of Traditions*, D. M. Fragaszy, S. Perry, Eds. (Cambridge University Press, 2003), pp. 33–55.

41. P. Todd, G. Miller, Exploring adaptive agency II: Simulating the evolution of associative learning. *From Animals to Animats,* 306–315 (1991).

42. C. R. Turner, T. J. H. Morgan, T. L. Griffiths, Environmental complexity and regularity shape the evolution of cognition. *Proc. Biol. Sci.* **291**, 20241524 (2024).

43. D. J. C. MacKay, *Information Theory, Inference and Learning Algorithms* (Cambridge University Press, 2003).
